# Dorso-ventral hippocampus neural assemblies reactivate during sleep following an aversive experience

**DOI:** 10.1101/2024.06.13.598918

**Authors:** JF Morici, A Silva, I Paiva-Lima, É Pronier, G Girardeau

**Affiliations:** Sorbonne Université, CNRS, Inserm, Center of Neuroscience NeuroSU, F-75005 Paris, France; Sorbonne Université, CNRS, Inserm, Institut de Biologie Paris-Seine, IBPS, F-75005 Paris, France

**Keywords:** hippocampus, emotional memory, sleep, neuronal assemblies

## Abstract

The integration of spatial and emotional components into episodic memory relies on coordinated interactions between hippocampal circuits and emotion-processing regions. While the dorsal hippocampus (dHPC) supports memory consolidation through sharp-wave ripple (SWR)-associated reactivation of spatial representations, it lacks direct connectivity with key emotional centers such as the amygdala. In contrast, the ventral hippocampus (vHPC) is anatomically and functionally embedded within the emotional network. How the dHPC and vHPC coordinate during sleep to support the processing of complex contextual and emotional experiences remains unclear. Here, we used simultaneous electrophysiological recordings from the dHPC and vHPC in rats performing a spatial alternation task under opposite emotional valence and during subsequent sleep. We show that dorso-ventral neuronal assemblies better disambiguate the between valence than isolated dorsal or ventral assemblies. During non-REM sleep, coordinated SWRs orchestrate assembly reactivation across the dorso-ventral hippocampal axis following both aversive and rewarding experiences; however, reactivation after aversive trials more closely mirrors the original neural patterns. This enhanced fidelity is driven by the heightened recruitment of shock-responsive neurons of the ventral hippocampus and spatial replay of the dorsal hippocampus into coordinated SWRs. Our findings identify a mechanism by which the hippocampus integrates the spatial and emotional dimensions of an experience during sleep, potentially routing information to the broader emotional memory network via synchronized SWRs.

## Introduction

Episodic memory is supported by the hippocampus^1^, a heterogeneous brain structure with distinct functional and anatomical properties along its dorsoventral axis^2,3^. The dorsal hippocampus (dHPC) encodes spatial representations of the environment through the activity of place cells, which code for specific locations called place fields^4^. During Non-REM sleep, these spatial representations are reactivated^5–7^, preferentially during fast oscillatory hippocampal events known as “sharp-wave ripples” (SWRs)^8^. Dorsal SWRs and the associated reactivation support spatial and contextual fear memory consolidation^9–11^. The integration of spatial information with other features of a lived experience, such as its emotional valence, involves a dialogue between the hippocampus and other structures of the emotional network, supported by the formation of cross-structure neural assemblies coordinated by SWRs^12–14^. There are joint reactivations between the hippocampus and the basolateral amygdala^15^ following an aversive spatial experience, and the processing of reward location involves a coordination between the dHPC and the ventral tegmental area^16^. Hippocampal spatial representations are modified by the emotional valence of the environment, a process known as remapping^17,18^ relying on the amygdala^19^. For historical and technical reasons, these studies have exclusively focused on the dorsal part of the hippocampus. However, the dHPC has no direct anatomical connections with the BLA and only marginal ones with the rest of the emotional network, including the mPFC, nucleus Accumbens and VTA^20,21^. This suggests that the inclusion of the dHPC in the emotional network for the integration of spatial and emotional information into complex memories requires an intermediate structure. The ventral hippocampus (vHPC) displays poorer spatial coding properties than the dHPC^3^, but plays a stronger role in emotional processing, particularly in anxiety-like behaviors^22–25^ and fear processing^26,27^. Unlike the dHPC, the vHPC has strong reciprocal anatomical and functional connections with the emotion-processing network, notably with the BLA^28–30^. The vHPC also displays SWRs, which are modulated by stress^31,32^. A marginal subset of ventral SWRs (vSWRs) are temporally coordinated with SWRs in the dHPC (dSWRs) in the absence of any prior learning or emotional experience ^33^. Ventral SWRs during sleep co-occur with fast oscillations in the BLA and mPFC and support coordinated reactivation between the vHPC and mPFC after fear conditioning^12^, but it is yet unclear whether vSWRs are associated with local vHPC^34^ and dorso-ventral hippocampal reactivation. Here, we hypothesized that during sleep, coordinated dorso-ventral SWRs support the reactivation of neuronal assemblies along the dorso-ventral axis of the hippocampus to process spatial and emotional information, potentially supporting their routing to structures selectively connected with the vHPC. Because the vHPC and connected emotional structures, like the BLA, encodes both positive and negative emotions^35–37^, we designed a behavioral task that allowed us to examine reactivation of a positive or negative experience triggering similar behaviors in the same spatial contexts. We performed simultaneous electrophysiological recording in the dHPC and vHPC in rats undergoing a spatial alternation task under rewarded or aversive conditions, and during the preceding and following sleep epochs. We identified dorsal, ventral, and dorsoventral neuronal assemblies in the hippocampus that represent the aversive and reward conditions of the task and investigated whether and how these assemblies differentially reactivate during subsequent NREM sleep and SWRs.

## Results

### Rats perform spatial alternation under rewarded and aversive conditions in the same context

To study the influence of emotional valence on dorsal-ventral hippocampal coordination, we developed a behavioral task in which animals learn a spatial alternation task on a linear track under two motivational contingencies associated with opposite valence. In the reward-motivated condition associated with a positive valence, the rats run back and forth on the linear track to obtain a small water reward at the end of each lap (“rewarded run”). The aversion-motivated condition, associated with a negative valence, was conducted on the same linear track and signaled by a single, constant light cue. Under this condition, animals run back and forth to avoid an eyelid shock (“aversive run”) delivered after 20-30 seconds of immobility (**Fig. 1**). The animals were first pre-trained to alternate sides in the reward condition. The aversive condition was then gradually introduced to preserve the mobility and alternation behavior (**Extended Data Fig. 1a**). The recordings started at plateau performance for both the reward and shock condition. Since the animals rapidly reach stable performance in the reward condition, plateau performance was defined as a drop in the number of shocks by at least 50% (**Fig. 1b**, lower panel). The number of rewarded laps and the number of aversive laps and shocks remained stable across all subsequent recording sessions (**Fig. 1b**, upper and middle panels; a lap is defined as one left-to-right or right-to-left run). Each recording day started with a baseline sleep and rest session in the homecage (sleep 1), followed by the aversive run and a second sleep session in the homecage (sleep 2). The animals then performed the rewarded run before the final sleep session in the homecage (sleep 3; **Fig. 1a**). The homecage sessions lasted 2 to 3 hours and the order of the reward and aversive conditions was switched every day. Animals explored homogeneously the linear track in both aversive and rewarded conditions (**Extended Data Fig. 1b**). The speed was globally lower in aversive laps compared to rewarded ones (**Extended Data Fig. 1c**), but the speed distributions across space were similar (**Extended Data Fig. 2a**). To control for behavioral differences across conditions, we used a speed threshold to divide the time on the linear track into active or quiet states. We did not find significant differences in occupancy in either of these states across space (**Extended Data Fig. 2b-c**) or in total time between conditions (**Extended Data Fig. 1d**). Subsequent neural analysis were restricted to active states, except for a control using quiet states.

**Figure 1:**
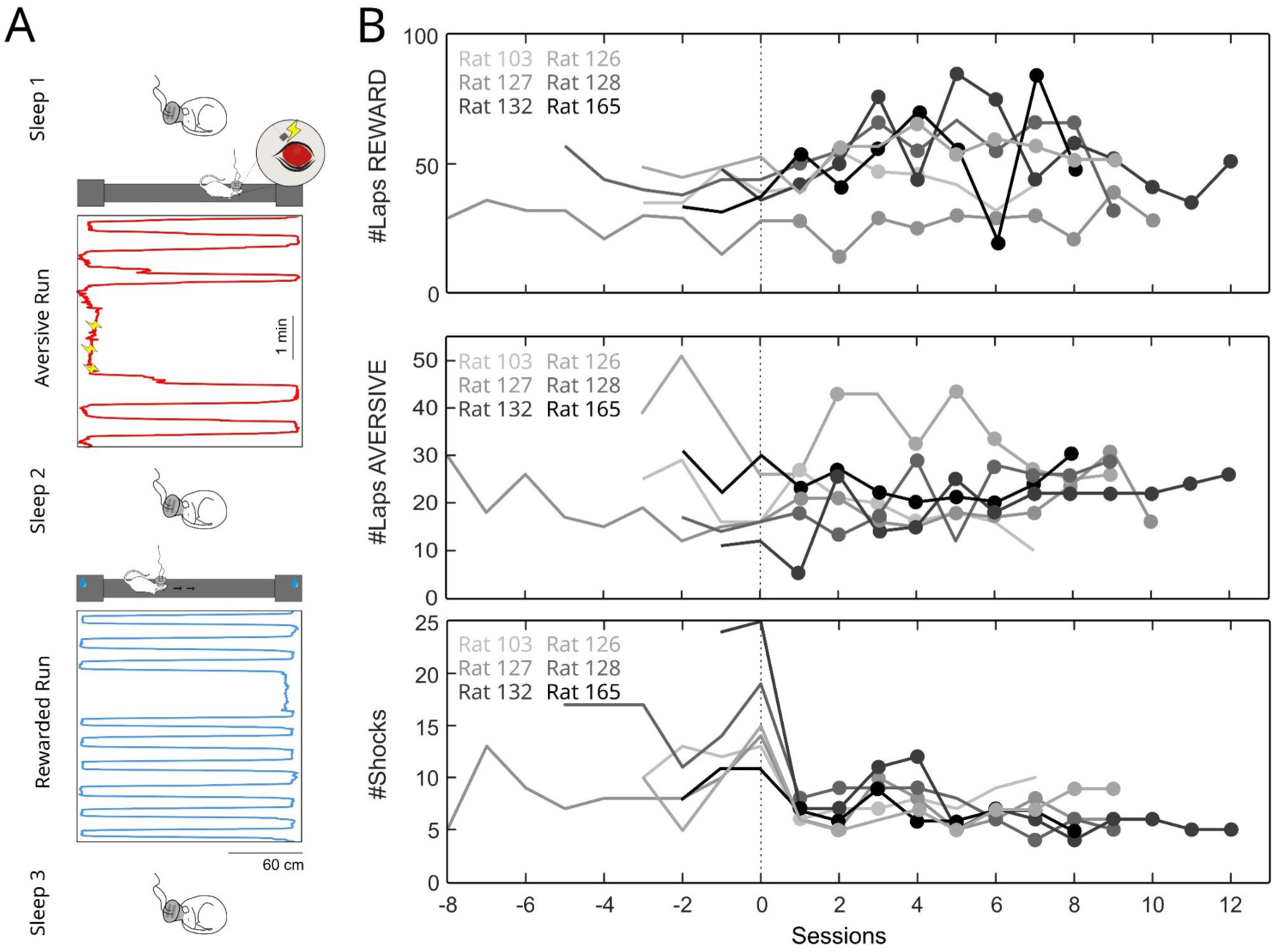
Rats perform aversive and reward-motivated alternation on a linear track. **A**, Example trajectories while rats run back and forth on a linear track to avoid eye-lid shocks (Aversive Run; red, yellow lightning indicate the shocks) or to receive a water drop (Reward Run; blue). Before and after each Run, animals were allowed to sleep in their home cage (Sleep 1-3). **B**, Number of rewarded laps, aversive laps and shocks during pre-training (-8 to 0) and during recordings days (1-12) at plateau performance starting as a 50% drop in the number of shocks received. Full dots indicate the recording days included in the analysis. (N_sessions_=47).

### Population and spatial coding in the vHPC better disambiguate the reward and aversive conditions

We recorded local field potentials (LFPs) and neuronal ensembles from the dorsal and ventral CA1 hippocampal regions with custom and commercial electrodes (**Fig. 2a-b ; Extended Data Fig. 3**) across a total of 47 recording days in 6 female Long-Evans rats. Each unit was classified as putative pyramidal cell (Pyr.) or interneuron (Int.) based on the spike waveform (see Methods, **Extended Data Fig. 4a-d**). In total, we identified 1,615 units (dHPC:22.79±2.22, vHPC:11.57±1.96, mean±sem units per session) across both regions: 544 units from the vHPC (Pyr: 461; Int: 83) and 1,071 from the dHPC (Pyr: 856; Int: 215). As expected, the firing rate was higher in putative interneurons compared to putative pyramidal cells in both regions, and putative pyramidal cells in the dHPC exhibited a higher burst index than vHPC putative pyramidal cells (**Extended Data Fig. 4e-f**, Connor & Gutnick, 1990). First, we analyzed the population-level representation in each hippocampal region during exploratory behavior in both aversive and rewarded runs. To do so, we applied principal component analysis (PCA) to the multi-neuronal activity from the dHPC and vHPC during each condition and calculated the Euclidean distance between the centroids of each condition’s coefficient cloud projected on PC 1 and 2. The Euclidean distance between the 2 conditions was significantly higher for the vHPC compared to the dHPC ( **Fig. 2c**), indicating that the population activity in the vHPC disambiguates better the rewarded and aversive conditions than the dHPC. Then we characterized place cell activity across the two regions and conditions. Spatial coding is known to be less precise in the vHPC compared to the dHPC^3,24^. Indeed, place cells made for 52% and 63% of putative pyramidal neurons in the dHPC, but only 17% and 29% in the vHPC in the aversive and rewarded conditions, respectively (**Extended Data Fig. 5a**). Place-fields were larger and peak firing rates lower in the vHPC compared to the dHPC, but neither these parameters nor the spatial information content differed between conditions in the dHPC or vHPC (**Extended Data Fig. 5b,c**). Finally, we assessed place cell remapping across conditions. We found that both structures remapped significantly between conditions compared to within conditions (**Fig. 2d-e**). When normalizing by the average within session spatial correlation, we found that remapping between conditions was higher in the vHPC than in the dHPC (**Fig. 2f**). Thus, the degraded spatial code found in the vHPC still contributes to the disambiguation between conditions, suggesting that place cells in the vHPC are more impacted by the change in contingency. Because the spatial context stays the same but the valence differs between conditions, this aligns with the preferential roles of the dHPC and vHPC in spatial and emotional processing, respectively^2,3^.

**Figure 2:**
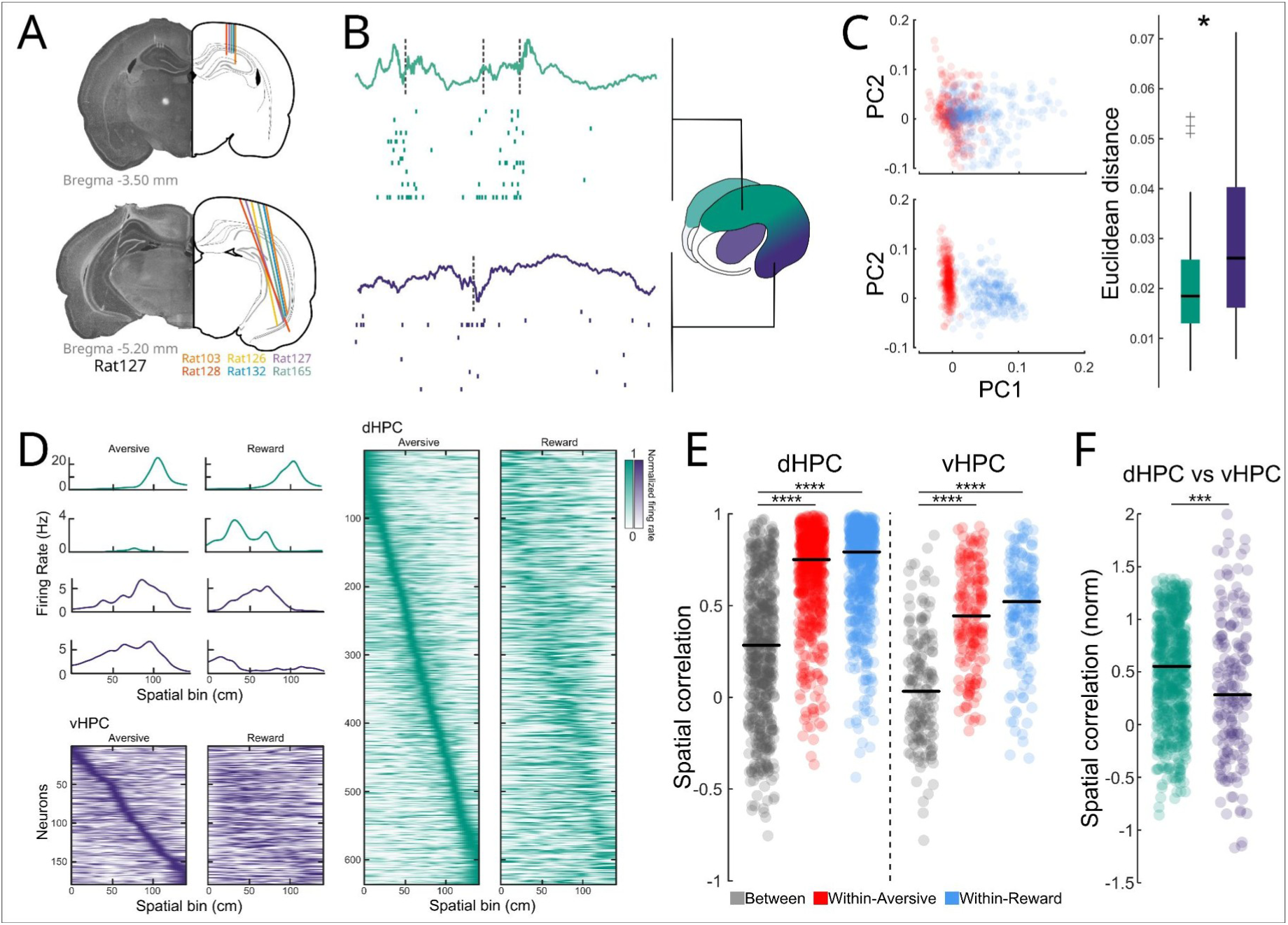
Ventral hippocampus population activity and spatial coding better distinguish aversive from reward sessions. **A**, Histological reconstruction of electrode placement in the dHPC and vHPC for all animals (1color/rat). **B**, Example LFP and spiking activity from the dorsal (green) and ventral (purple) hippocampus during NREM sleep. Dotted lines indicate SWRs. **C**, Example of PCA-projected population spiking activity during exploration behavior in the rewarded (blue) and aversive (red) runs for the dorsal (top) and ventral (bottom) single-units for one recording session (left panels). Comparison of Euclidean distance between centroids of reward and aversive PCA-projected clouds obtained with the dorsal and ventral hippocampal population activity for all recording sessions (right panel; two-sided Mann-Whitney test, *p =0.012, N_dorsal_= 37, N_ventral_=35, black line: median). **D**, Top left : Pyramidal cells identified as place cells in at least one session. Place-fields of four examples place cells of the dHPC (green) and vHPC (purple) in the aversive and reward conditions. Bottom left and right : All cells identified as place cells in at least one session, sorted according to the position of their peak response during the aversive run in the dHPC (green, right) and vHPC (purple). **E**, Spatial correlation of tuning curves within reward (blue) or aversive (red) sessions, and between conditions (gray), for dorsal (left) and ventral (right) place cells (Friedman test, ****p < 0.0001; Tukey–Kramer post hoc test, ****p < 0.0001). **F**, Normalized spatial correlation between reward and aversive sessions for dorsal (green) and ventral (purple) place cells (unpaired Wilcoxon test, ***p = 0.0003).

### The shock differentially modulates neuronal responses in the dorsal vs. ventral hippocampus

Both the dorsal and ventral hippocampus can respond to aversive stimuli during fear conditioning^26,38,39^. Here, the shock was not associated to a cue or a context, but to a specific behavior (immobility), and meant to promote a structured avoidance behavior (i.e. alternation) in the absence of a defined safe location. This creates a threatening condition in the otherwise stable context. To check for shock-responsiveness in our task, we constructed firing curves for putative pyramidal cells in both the dorsal and ventral hippocampus (**Fig. 3a**). We found that a significantly higher proportion of neurons in the dHPC exhibited an increase in their response during the shock (**Fig. 3b**). However, the shock systematically increases the speed of the animal (**Fig. 3c**), and speed is known to influence the firing of dorsal and ventral hippocampal neurons, albeit to a different extent : dorsal neurons are more speed-modulated than ventral neurons^40^. In our data, the proportion of speed-modulated neurons within the vHPC shock-responsive neurons was the same as for vHPC non shock-responsive neurons. In the dHPC, the shock-responsive neurons were slightly enriched in speed-modulated neurons compared to the non shock-responsive neurons (**Extended Data Fig. 6a-c**). To account for the bias in shock response due to the increase in speed, we decorrelated speed from the neuronal response (see Methods). While dorsal units exhibited a significant decrease in the magnitude of shock-response upon decorrelation, ventral shock-responsive units showed no significant change. Once controlled for speed, the vHPC shows a stronger response to the shock than the dHPC (**Fig. 3d-e**). Contrary to the dHPC, shock cells in the vHPC do not increase their activity at transitions from quiet to active behavior in the absence of shock. In shock-responsive neurons, the maximal shock response significantly correlated with the magnitude of speed modulation for the dHPC neurons but not for the vHPC neurons. (**Extended Data Fig.6d-e**). We also investigated the link between shock cells and place cells. In the dHPC or vHPC, the proportion of shock-responsive neurons is not significantly different among place-cells vs. non-place cells. The place-fields of shock-responsive place cells were not closer to the location of the first shock compared to the place-fields of non shock-responsive place-cells, indicating that the shock does not turn nearby place-cells into shock-responsive neurons. Dorsal shock-responsive place cells exhibited lower spatial information values than pure place cells, whereas this effect was absent in the ventral hippocampus where spatial information is already significantly lower (**Extended Data Fig. 6f-h**). In the rewarded condition, we found a similar proportion of reward-responsive cells in the dHPC and vHPC (**Fig 3f-g**). Because the reward is consumed during immobility (**Fig. 3h**), speed-decorrelation does not affect reward-responsiveness in either structure, but the responses are higher in the dHPC (**Fig. 3i,j**). Altogether, these results indicate that both the dHPC and vHPC respond to shock and reward. In the dHPC, shock-responsiveness is strongly influenced by speed, and to a lesser extent space, whereas the vHPC exhibits strong shock responsiveness largely independent from speed- and place-modulation.

**Figure 3:**
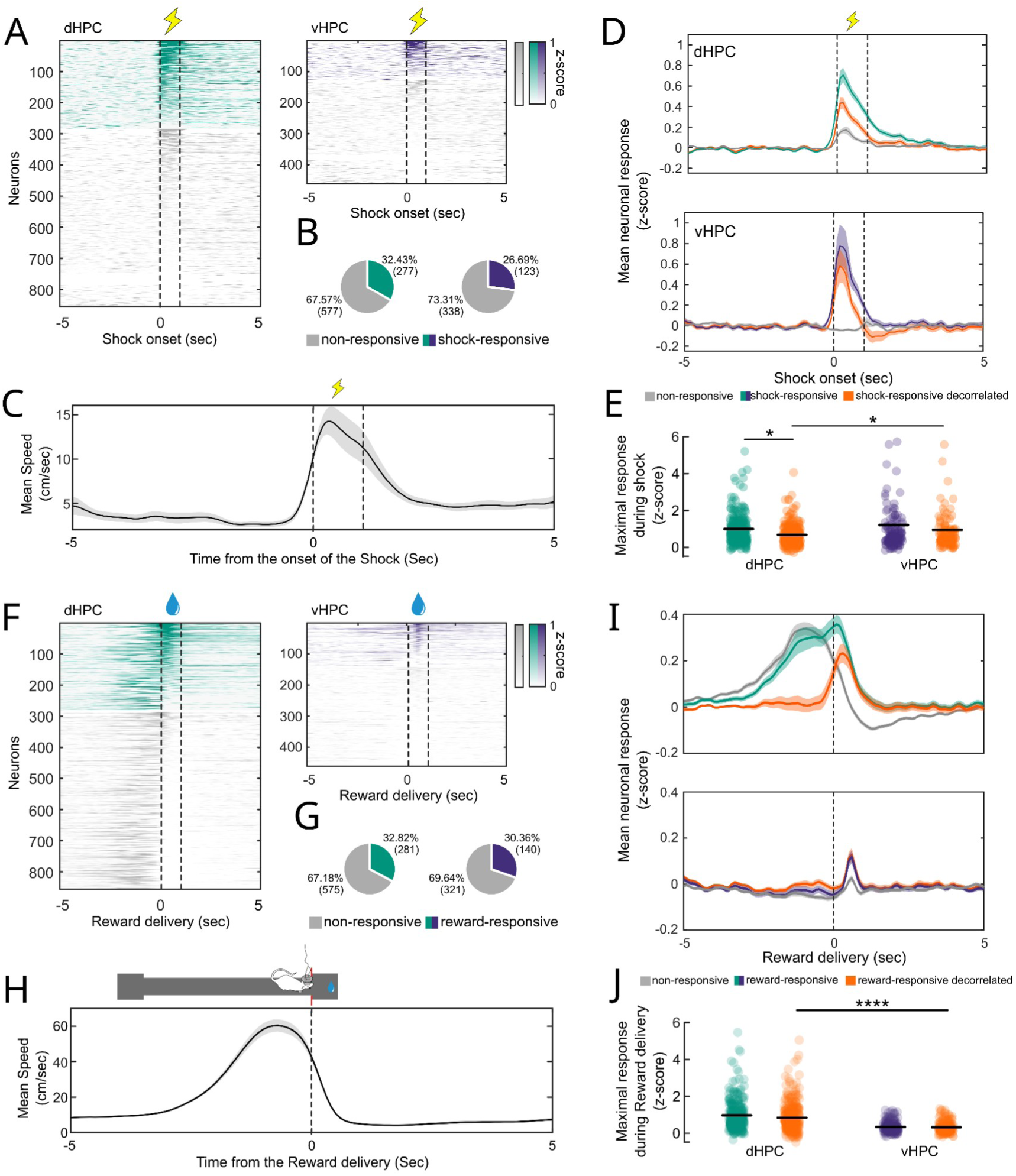
Shock-responsiveness is higher in the ventral than in the dorsal hippocampus. **A,** Normalized peri-shock responses for all dHPC and vHPC neurons. **B,** Percentage of shock-responsive neurons in the dHPC and vHPC (Chi^2^, *p=0.03). **C**, Average speed centered on shock onset for all aversive sessions in all animals (N_sessions_=47, N_shocks_=4-13). **D**, Mean z-scored response to shocks for shock-responsive neurons (green/purple) and non-responsive neurons (gray) in the dHPC (top) and vHPC (bottom). When decorrelated from speed (orange), the response of shock-responsive neurons decreases significantly more in the dHPC than in the vHPC. **E**, Maximal response during shock for shock-responsive (green/purple) and shock-responsive decorrelated activity (orange; paired Two-way ANOVA, p_structure_=0.051, F=3.8, **p_speed_<0.004, F=8.24, p_interaction_=0.59, F=0.29. Tukey-Kramer post-hoc test with Bonferroni correction, *p=0.01, n_dHPC_ = 277, n_vHPC_=123; black line: mean). **F**, Normalized peri-reward responses for all dHPC (green) and vHPC (purple) neurons. **G**, Percentage of shock-responsive neurons in the dHPC and vHPC (Chi^2^, p=0.36) . **H**, Average speed centered on reward trigger for all reward sessions in all animals (N_sessions_=47, N_rewards_=9-86). I, Mean z-scored response to valves opening for reward-responsive neurons (green/purple) and non-responsive neurons (gray) in the dHPC (top) and vHPC (bottom). **J**, Maximal response during reward consumption for reward-responsive (green/purple) and shock-responsive decorrelated activity (orange; paired Two-way ANOVA, ****p_structure_ <0.00001, F=120.3, p_speed_ =0.23, F=1.41, p _interaction_=0.36, F=0.85. Tukey-Kramer post-hoc test with Bonferroni correction, ****p<0.00001, N_dHPC_ = 281, N_vHPC_ =140).

### Dorso-ventral hippocampal joint assemblies better disambiguate the reward and aversive conditions

Functional communication between structures relies on the formation of neuronal assemblies composed of members distributed across these regions^14,41,42^. We examined whether the dorsal and ventral hippocampus could bind spatial and emotional information via the formation of dorso-ventral neuronal assemblies. The assemblies were detected during the aversive or the rewarded condition using a PCA-ICA method^43^, **Extended Data Fig. 7**) on all simultaneously recorded pyramidal neurons in the dorsal and ventral hippocampus. The detection was restricted to moments of active behavior (**Extended Data Fig 2c**). For each of the 446 detected assemblies, we applied a threshold to the ICA weight vectors to determine the neurons that were significantly contributing to the assembly ( **Fig. 4a**). We differentiated 277 assemblies with only dorsal hippocampus contributing members (131 aversive and 146 rewarded), 62 assemblies with only ventral hippocampus members (37 aversive and 25 rewarded), and 107 assemblies with contributing members from both the dorsal and ventral hippocampus. We called these assemblies “joint assemblies” (54 aversive and 53 rewarded, 23,99% of detected assemblies). We did not detect differences in the number or proportion (**Extended Data Fig. 8**) of members across assembly types (dorsal, ventral, or joint). Most dorsal and ventral members of joint assemblies were unique to those assemblies, rather than shared with purely region-specific assemblies (aversive dHPC, 66,66%, N=120, reward dHPC, 60,23%; aversive vHPC, 82,66%, N=143; reward vHPC, 88,30%, N=166). To compare the overlap between assemblies detected either in the aversive or rewarded condition on the same set of neurons, we computed the similarity index across pairs of dorsal, ventral, or joint assemblies and identified the pairs that were significantly similar between the two conditions (**Fig. 4b**). The percentage of assemblies that overlapped between the two conditions was significantly lower for the joint assemblies compared to the dorsal or ventral assemblies (**Fig. 4c**). Hence, while the global activity in the ventral hippocampus alone disambiguates better the valence of the context than the dorsal hippocampus (**Fig. 2c, f**), at the level of assemblies specifically involved in each condition, representations associating both the dorsal and ventral hippocampus lead to a better disambiguation of the rewarded vs. aversive experience in the same context.

**Figure 4:**
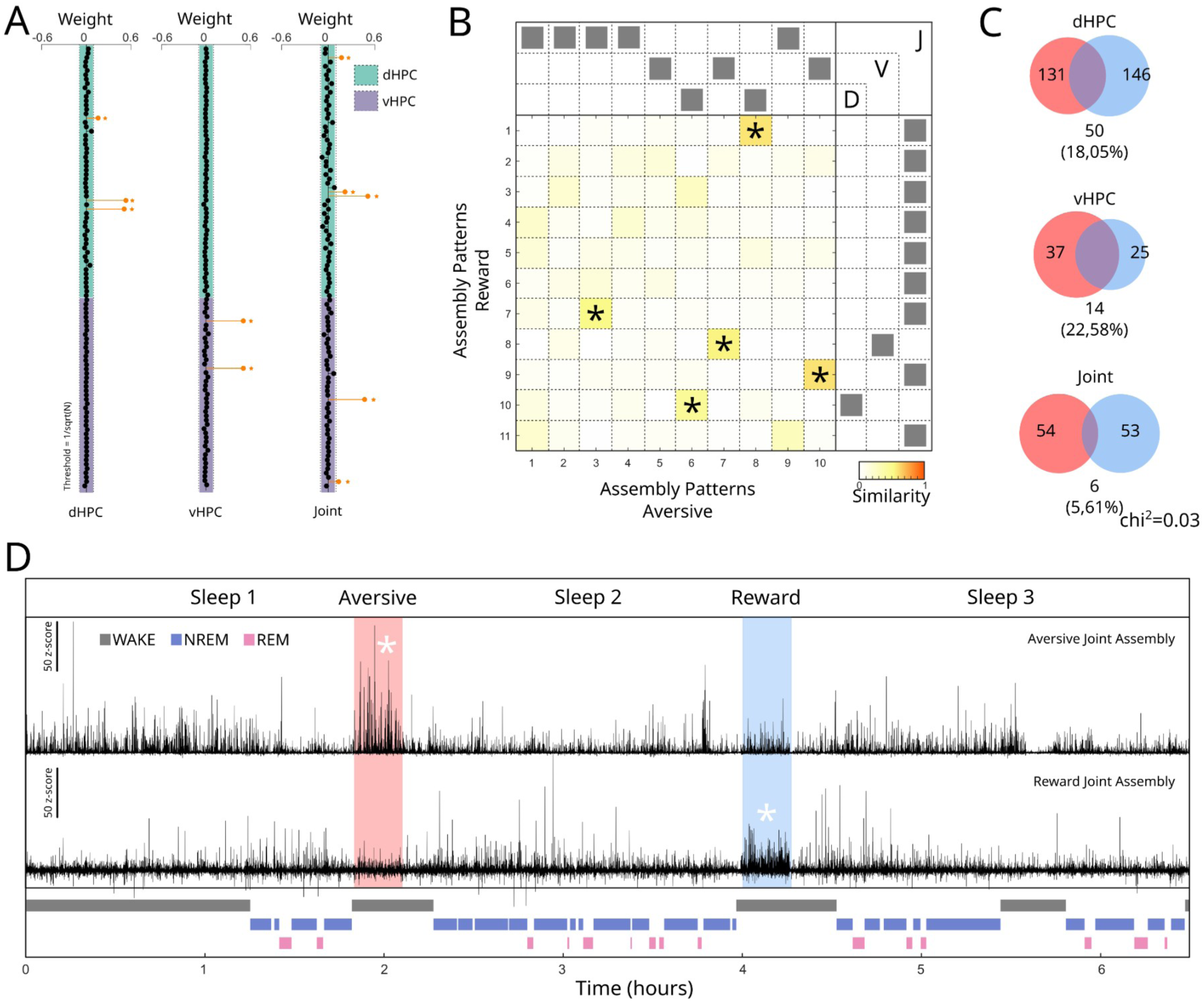
Assembly detection during Aversive and Rewarded runs. **A**, Examples of a dHPC (left), vHPC (middle) and joint (right) assembly from a representative session with assembly members (orange lollipops) in the dHPC, vHPC or both, respectively. **B**, Example of an Assembly Similarity Index matrix for dHPC, vHPC and joint assemblies. The index was calculated as the dot product of all assembly pairs in the session. Two assemblies were considered significantly similar (*) if the Similarity index was above the 99.9 percentile of a surrogate distribution. **C**, Venn diagrams of significantly similar aversive (red) and rewarded (blue) assemblies for dHPC, vHPC and joint assemblies. Joint assemblies showed the lowest overlapping between conditions (Chi^2^, *p=0,03). **D**, Examples of assembly activity strength during the whole recording session for a joint aversive assembly (red) and a joint rewarded assembly (blue).

### Dorsal, ventral and joint hippocampal assemblies reactivate during sleep after the rewarded and aversive conditions

During sleep after an experience, the neural patterns of exploration show an increased activation compared to pre-experience sleep. This reactivation phenomenon occurs within the dorsal hippocampus^5,7^ and between the dorsal hippocampus and structures related to valence processing^3,15,16,44^. The extent to which reactivation occurs in the vHPC is yet unclear^45^, and it is unknown how the vHPC coordinates with the dHPC during sleep following an emotional experience. To address this question, for each joint assembly detected during the aversive or reward experience, we calculated the activation strength during the rest of the recording session, including REM and NREM sleep in the homecage (**Fig. 4D**). We used a threshold to identify activation peaks and calculated the activation peak rate per minute. This quantitative measure indicates the frequency at which the assembly appears during NREM sleep before and after the rewarded or aversive exploration (**Fig. 5a**). Both aversive and reward joint assemblies exhibited higher activation rates during post-sleep compared to pre-sleep periods, establishing the existence of joint dorso-ventral hippocampal reactivation (**Fig. 5b**). The pre-to-post sleep difference diminishes over the course of the sleep session (0-20 minutes vs 20-40 minutes of NREM sleep; **Fig. 5b**), consistent with previous observations that reactivation rapidly declines over the first hour of NREM sleep^12,15^. The same result was observed for dorsal and ventral assemblies (**Extended data Fig. 9**). This was not due to concatenating fragmented NREM bouts into a single block (see Methods), because the condition did not modify the general structure of the following sleep (REM and NREM bouts # and durations; **Extended Data Fig. 10 and 11**), and NREM assembly reactivation also showed a significant decay in real time elapsed from the beginning of the first NREM epoch (**Extended Data Fig. 12**). No reactivations were found during REM sleep (**Extended data Fig. 13a**). Thus, our results show that NREM sleep reactivation after an emotional experience extends beyond the dorsal hippocampus and includes reactivation in the ventral hippocampus as well as coordinated reactivation across the full hippocampal axis after both rewarded and aversive experiences.

**Figure 5:**
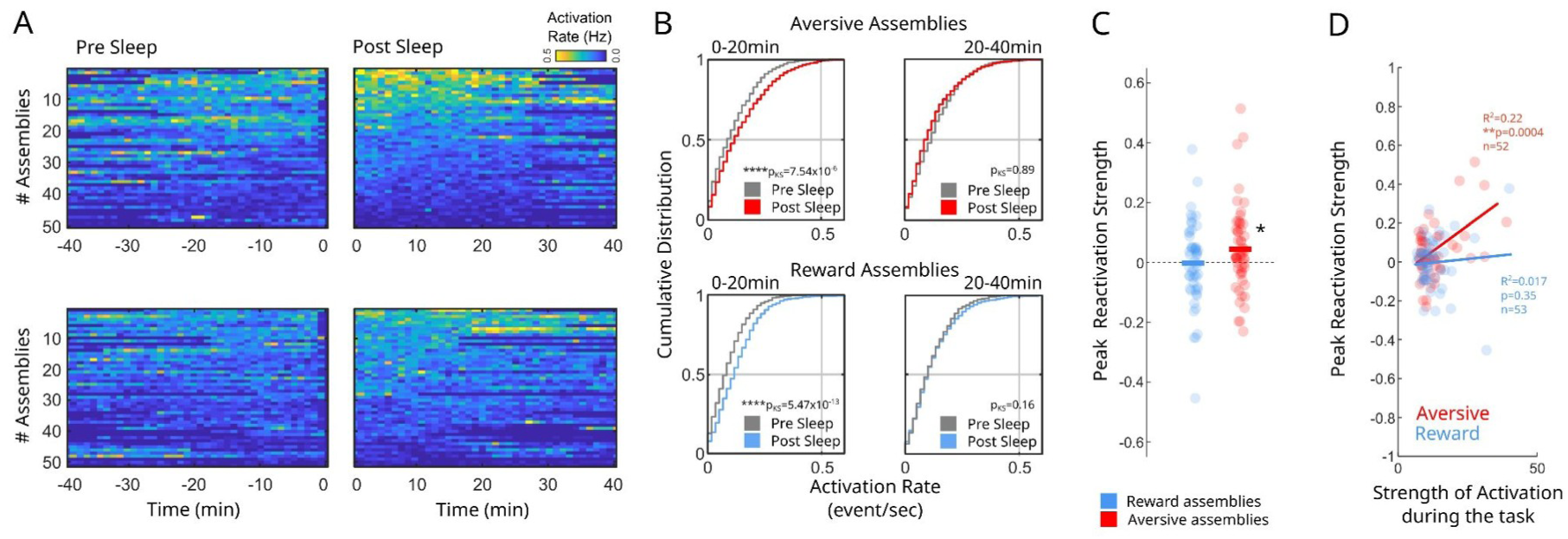
Joint assemblies reactivation during NREM sleep. **A**, Assembly peak activation rate across time during pre- and post-sleep sessions for all aversive (top) and rewarded (bottom) assemblies (N_aversive_=50, N_rewarded_=51). **B**, Cumulative distributions of peak activation rates including pre-sleep (in gray) and and post-sleep reactivation rates for aversive (red) and rewarded (blue) assemblies Note the significant increase in activation rate in post vs. pre-sleep for both types of assemblies in the first 20 minutes of NREM sleep. (Kolmogórov-Smirnov test, ****p<0,0001). **C**, Peak Reactivation Strength for aversive and reward assemblies, significantly different from zero for aversive joint assemblies only, . (one-sided Mann-Whitney test against zero, *p=0,014; Mann-Whitney test, p=0.08; N_aversive_=54, N_reward_=53). **D**, Joint aversive assemblies’ Reactivation strength correlates with their activation during the task. (Linear regression; Aversive, R^2^=0.22, **p_slope_<0.001; Reward, R^2^=0.017, p_slope_=0.35).

### Aversive joint assemblies reflect more accurately the awake experience

Next, we investigated potential differences between the joint dorso-ventral assemblies reactivation after a rewarded vs. an aversive experience. We calculated the average activation strength across peaks during NREM sleep before and after exploration. This qualitative measure reflects the similarity in co-activation patterns of members between the assemblies activated during NREM sleep and the “original” assemblies detected during the aversive or rewarded experience. Then we computed the pre-to-post-sleep ratio between these values to obtain the peak reactivation strength (see Methods). A positive peak reactivation strength indicates that assemblies activated in sleep after the experience show a higher fidelity to the experience assemblies compared to assemblies in sleep preceding it. We observed that the peak reactivation strength was significantly different from zero for the joint aversive assemblies but not for the reward joint assemblies (**Fig. 5C**), nor for the aversive or reward dHPC or vHPC assemblies (**Extended data Fig. 14a**). The peak reactivation strength was not different from zero for any assembly type during REM sleep (**Extended data Fig. 13b**). Thus, when reactivated during post-experience NREM sleep, the joint aversive assemblies are more similar to the original assemblies that emerged during the aversive experience compared to pre-experience NREM assemblies. Finally, we found that the peak reactivation strength of joint aversive assemblies positively correlated with the activation strength of the assemblies during the exploration of the aversive condition ( **Fig. 5d**). This correlation was selective for joint aversive assemblies, and not significant for either joint reward assemblies or dHPC/vHPC assemblies in either condition (**Fig. 5d**, **Extended data Fig. 14b**). This indicates that the increase in peak activation strength of the aversive joint assemblies during NREM sleep was initiated during the preceding exploration of the track. Indeed, joint-assembly peak reactivation strength showed a positive correlation with the percentage of active exploration during aversive run sessions, but not in the reward condition, where movement itself has no behavioral relevance. Peak reactivation strength was not influenced by the number of shocks or rewards (**Extended Data Fig. 15**). This suggests that behavioral engagement in the aversive session—where animals avoid shocks by actively moving through space—influences the strength of reactivation during subsequent sleep. Altogether, our results show that spatial exploration under rewarded or aversive conditions recruits assemblies spanning both the dorsal and ventral hippocampus. These assemblies will be later reactivated during NREM sleep, with joint aversive assemblies showing a strong fidelity to the assemblies that emerged during the experience.

### Coordinated SWR following the aversive condition are associated with increased dorsal sequence replay

Dorsal hippocampus reactivations have consistently been found to co-occur with sharp-wave ripples (SWRs)^46^. SWRs were shown to mediate the synchronization of the dHPC activity with distant structures^13,15,16^. While a subset of SWRs coordinate between the dHPC and vHPC during baseline NREM sleep^33^, it is yet unknown how spatial exploration associated with an emotional valence affects vHPC SWRs and their coordination with dHPC SWRs during subsequent sleep. To detect SWRs in the dorsal (dSWRs) and ventral (vSWRs) hippocampus, we used the oscillatory component (ripple) of the SWRs by thresholding the normalized square root of the filtered local-field potential recorded in the CA1 pyramidal layer (**Fig. 6a-b**). As previously established, we found that dSWR rates were significantly higher than vSWR rates (**Fig. 6c)**^33^. However, the vSWR rate did not increase during sleep following either the rewarded or the aversive session, as opposed to dSWRs (**Extended data Fig. 16a**). Ripples tend to occur in bursts, as shown by the shape of the autocorrelograms (**Extended data Fig. 16b**). We did not observe changes in the “burstiness”, amplitude or duration of dSWRs or vSWRs neither after the aversive nor the rewarded experience (**Extended data Fig. 16c-e**).

**Figure 6:**
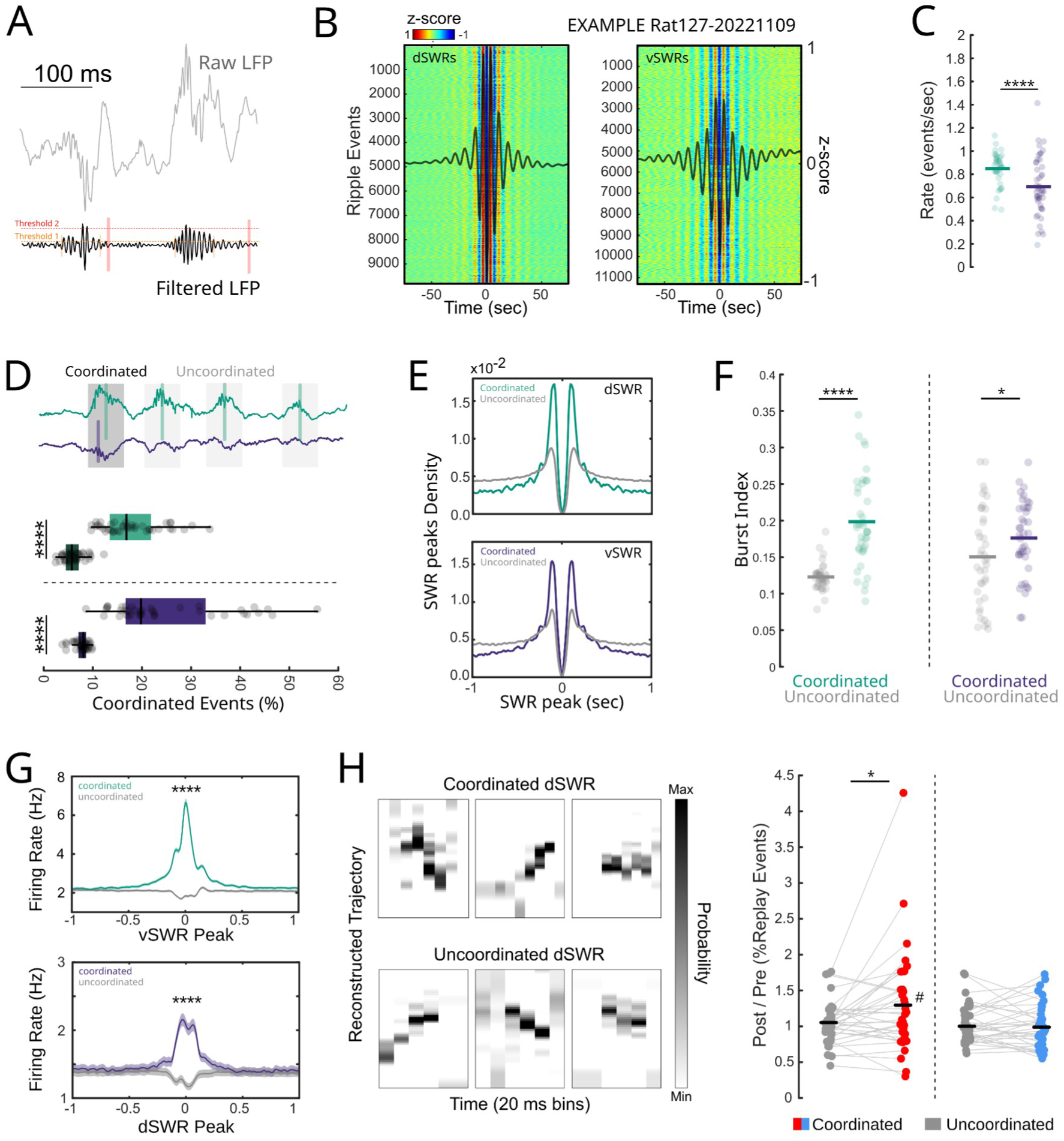
Ripple coordination across dorsal-ventral hippocampal axis preferentially sustains replay of spatial sequences. **A,** Example raw and filtered LFP of the dorsal hippocampus during NREM sleep. Ripples in both regions were detected by thresholding the normalized squared local-field potential signal filtered between 100Hz and 250Hz. **B**, Colormap showing z-scored signal surrounding the peak of each detected dorsal (left) and ventral ripple (right). **C**, Ripple rate calculated during NREM sleep in the dorsal (green) and ventral (purple) hippocampus (two-sided Mann-Whitney test, ****p=0,0007, N_dHPC_=40, N_vHPC_=47). **D**, Coordinated events were defined as a pair of dorsal-ventral ripples that occurred within 100ms of each other. Example dHPC and vHPC LFP traces showing 1 coordinated ripple (Top). The percentage of coordinated events in the dHPC (green) and vHPC (purple) was different from chance level (black; two-sided Mann-Whitney test, ****p<0.0001, N=40, box edges are 25th and 75th percentile, line represent median). **E**, Average autocorrelogram of ripple peaks for coordinated (green/purple) and uncoordinated (gray) ripples in the dHPC (top) and vHPC (bottom). **F**, Burst Index of coordinated and uncoordinated ripples from the dorsal and ventral hippocampus. (two-sided Mann-Whitney test, *p<0.05, ****p<0.0001, N=40). **G**, Response of dorsal (top) and ventral (bottom) HPC neurons to coordinated (green/purple) and uncoordinated (gray) ripples (paired Wilcoxon test, ****p<0.00001, N_dHPC_=856, N_vHPC_=461). **H**, Left panel, Examples of significant dorsal replay events during dorsal ripples occurring within coordinated (top) and uncoordinated (bottom) ripple events. Right panel, Ratio of the pre/post percentage of significant replay events for coordinated (red or blue) and uncoordinated (gray) dorsal ripples for aversive (red) and rewarded (blue) sessions (Paired Wilcoxon test, *p_aversive_=0.02, p_reward_=0.77, N=40. One-sample Wilcoxon test against one, ^#^p=0.018).

Next, we explored the fine temporal coordination between dorsal and ventral ripples. On the cross-correlation between dorsal and ventral ripples, the central peak was shifted to positive time bins, suggesting that vSWRs tend to occur after dSWRs (**Extended Data Fig. 17a**). We defined coordinated events as events during which a vSWR occurred within +/-100 ms of a dorsal one. Both dorsal and ventral ripples showed a percentage of coordinated events significantly higher than chance ( **Fig. 6d**, Median, dHPC: 16.83%, vHPC: 19,81%). Surprisingly, neither the proportion of coordinated events nor their directionality (dHPC vs vHPC leading) changed after either the rewarded or the aversive condition compared to baseline (**Extended data Fig. 17b-d**), suggesting that changes in coordination occur at a finer scale. The amplitude of coordinated dorsal and ventral ripples was higher than the amplitude of uncoordinated ripples (**Extended data Fig. 17e**), as reported previously^33^, and coordinated ripples occur more in bursts than uncoordinated ones (**Fig. 6e-f**). The recruitment of single dHPC and vHPC by units by dorsal or ventral ripples, respectively, does not differ between coordinated and uncoordinated ripples (**Extended data Fig. 17f-h**). Dorsal neurons and ventral neurons do not respond to uncoordinated vSWR and dSWR, respectively, but both dorsal and ventral neurons respond during coordinated SWRs of the opposite pole (**Fig. 6g**). This suggests that coordinated events represent a privileged time window for cross-hippocampal neuronal communication.

Dorsal ripples are known to support the reactivation of spatial sequences^5,6^. Because we hypothesized that coordinated events provide a time window in which dorsal spatial information and ventral emotional information are jointly engaged, we investigated whether the spatial content in the dHPC differed between coordinated and uncoordinated SWRs. We detected sequence replay using a Bayesian decoding method and found that 8.81 ± 0.63% of dorsal ripples contained significant replay events^49^. We then separated significant replay events occurring during coordinated versus uncoordinated SWRs. The percentage of significant replay was significantly increased during coordinated SWRs following the aversive condition, but not the reward condition (**Fig. 6h)**. These results were replicated using rank-order correlation as an alternative detection method (**Extended Data Fig.18**). Together, these results indicate that coordinated SWRs are enriched in relevant spatial content after an aversive experience.

### Coordinated SWRs support stronger joint aversive assembly reactivation through increased recruitment of vHPC shock-responsive neurons

We hypothesized that coordinated dorso-ventral SWRs support the reactivation of joint assemblies. We showed that the peak reactivation strength for joint aversive assemblies is significantly different from zero and significantly different from the peak reactivation strength of rewarded assemblies during coordinated SWRs, but not during uncoordinated SWRs (**Fig. 7a**). This effect was not due to merging uncoordinated vSWRs and dSWRs (**Extended data Fig. 19**) and was specific to joint assemblies (**Extended data Fig. 20a-b**). To rule out the possibility that the absence of increased peak activation for reward joint assemblies was due to excluding quiet awake periods—when reward consumption and synchronous activity could occur—we repeated the analysis using assemblies detected during quiet periods only and recalculated Peak Reactivation Strength during coordinated and uncoordinated NREM ripples. Peak reactivation strength was not different from zero either aversive or reward “quiet” joint assemblies (**Extended data Fig. 20c**). These results indicate that dorso-ventral coordinated ripples constitute a permissive time window for the reinstatement of the representation of an aversive environment by the joint neural activity of the dorsal and ventral hippocampus.

**Figure 7:**
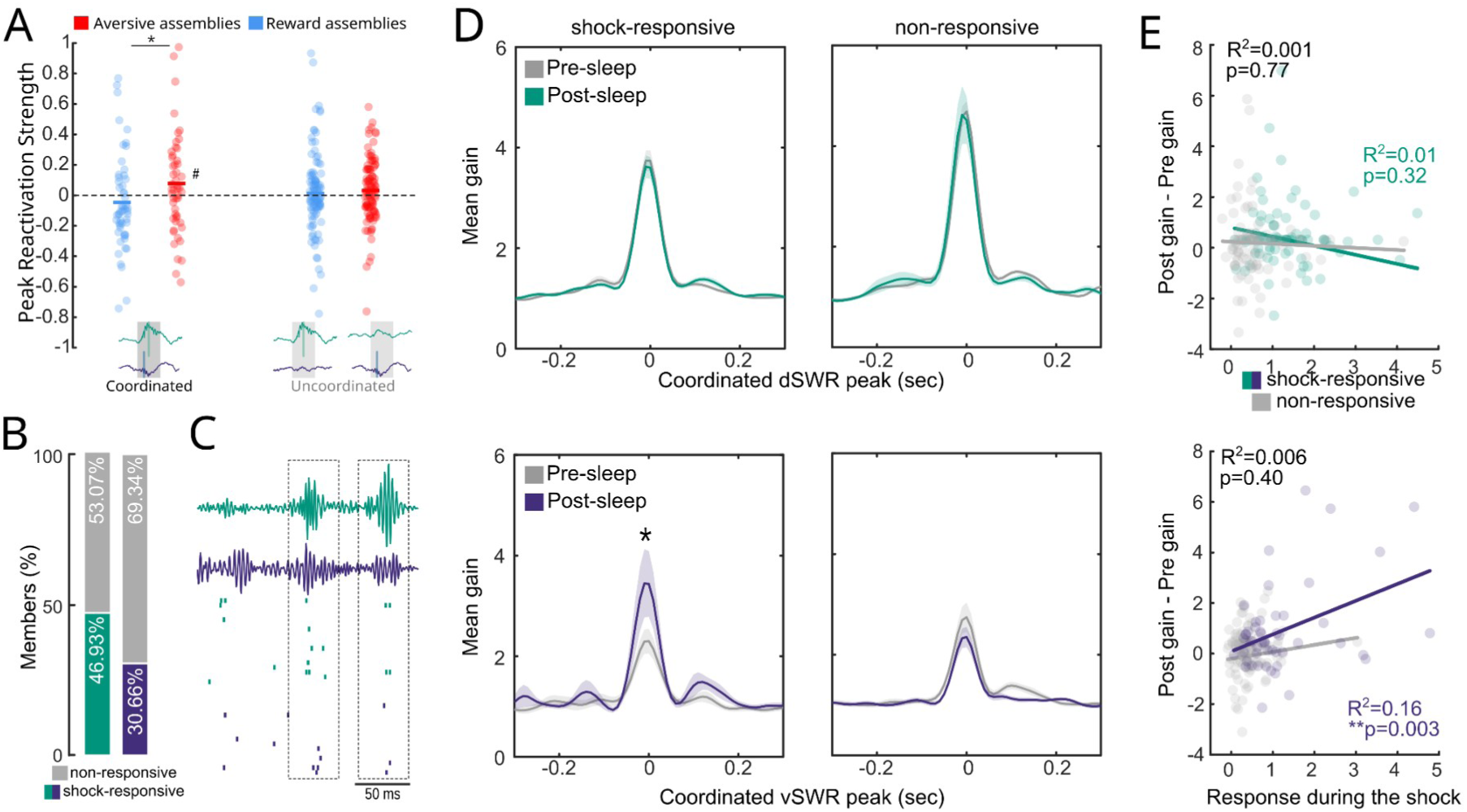
Shock-responsive ventral members increase their gain during dorsal-ventral coordinated ripples. **A**, Average peak reactivation strength for joint aversive and reward assemblies during dorsal-ventral coordinated ripples and dorsal or ventral uncoordinated ripples (non-paired Two-way ANOVA, p_interaction_=0.11, F=2.45, *p_assembly_type_=0.02, F=5.07, p_ripple_type_=0.92, F=0.01. Tukey-Kramer post-hoc test with Bonferroni correction, *p_coordinated_=0.024, p_uncoordinated_=0.25). Only joint-aversive assemblies showed a significant increase against zero. (one-sided t test, ^#^p=0.04, N_aversive_=54, N_reward_=53). **B**, Percentage of shock-responsive neurons among dHPC (green) and vHPC (blue) members of joint aversive assemblies (dHPC:46.93%; vHPC: 30.66%). **C**, Example of two coordinated dorsal-ventral ripple events (dashed box), (top) and associated single unit activity (bottom). **D**, Average gain in firing rate around coordinated ripple onset during NREM sleep preceding (grey) and following (green/blue) avesrive runs. (one-sided Mann-Whitney test. dHPC, p_responsive_=0.86, N=84, p_non-responsive_=0.99, N=95. vHPC, *p_responsive_=0.039 , N=53, p_non-responsive_=0.99, N=120). **E**, Linear correlation between the maximal response during the shock delivery and the change in gain between pre- and post-sleep for coordinated ripple events. (Linear regression, dHPC: Shock-responsive, R^2^=0.01, p_slope_=0.32, Non-responsive, R^2^=0.001, p_slope_=0.77; vHPC: Shock-responsive, R^2^=0.16, p_slope_=0.003, Non-responsive, R^2^=0.006, p_slope_=0.40).

We sought to identify the origin of the specific increase in peak reactivation strength for joint aversive assemblies. In particular, we hypothesized that vHPC shock-responsive neurons are key in this phenomenon. However, neither dorsal or ventral members of the joint aversive assemblies were enriched in shock-responsive units compared to the whole population of units from each region ( **Fig. 7b**, Chi^2^; vHPC: X^2^=0.71, p=0.40; dHPC: X^2^=0.13, p=0.72). Hippocampal neurons are globally massively recruited during SWRs, but experience modulates this recruitment^47^. We assessed the gain (i.e. the normalized increase in activity) of each vHPC or dHPC member of the joint assemblies during coordinated SWRs before vs. after the aversive or rewarded experience. Additionally, we distinguished between shock-responsive members and non shock-responsive members (**Fig. 7b,c**). In the dorsal hippocampus, neither the shock-responsive nor the non-responsive members showed an increase in gain during post-NREM coordinated SWRs compared to pre-NREM SWRs (**Fig. 7d**, upper panel). However, in the ventral hippocampus, we found that shock-responsive members, but not non-responsive ones, significantly increased their activity during post-NREM coordinated SWRs (**Fig. 7d**, lower panel). This response was specific to coordinated SWRs (**Extended data Fig. 21**) and shock-responsive neurons, as there were no significant changes for reward-responsive neurons during coordinated or uncoordinated SWRs (**Extended data Fig. 22**). Shock- and reward-responsive members of pure dHPC or vHPC assemblies did not increase gain in coordinated or uncoordinated SWRs either (**Extended data Fig. 23)**. Finally, we found a positive linear correlation between the change in activity gain during SWRs from pre- to post-sleep and the level of activation during the shock experience selectively for ventral shock-responsive neurons (**Fig. 7e**). Altogether, these results suggest that the enhanced fidelity of joint reactivation after aversive experience is due to a selective increase in the participation of shock vHPC responsive neurons to the coordinated assemblies during SWRs, originating in the intensity of their recruitment during the awake experience.

## Discussion

In this study we detected dorsal, ventral and dorso-ventral hippocampal neural assemblies during a similar spatial experience under two opposite valence conditions. We then tracked them during the subsequent epochs to investigate the sleep-dependent reactivation mechanisms known to underlie memory consolidation within the dHPC and across the dHPC and other structures^15,16,48^. Our findings establish for the first time the existence of dorso-ventral hippocampal coordinated reactivation during Non-REM sleep and provide further support for the independent reactivation of the vHPC^45^. In our task, the spatial context is stable and only the emotional valence changes. Given the respective roles of the dorsal and ventral hippocampus in spatial and emotional processing^4,23,24^, and the corresponding dynamics of their neuronal assemblies—spatially tuned and pre-configured in the dHPC^7,49^, versus emerging during emotional experience in the vHPC^12,38,50^—we expected dorsal assemblies to remain more consistent across the two conditions than ventral ones. Surprisingly, joint assemblies, rather than pure dorsal or ventral assemblies, better disambiguate the two conditions. The higher remapping of place-cells between conditions in the vHPC compared to the dHPC contributes to the disambiguation. We hypothesize that dorso-ventral coordination during acquisition and sleep-dependent processing allows to functionally bind the stable spatial representation of the dorsal hippocampus with the bimodal emotional aspect of the experience processed in the vHPC to compose an integrated, refined representation embedding all contextual parameters.

Both reward and aversive assembly show reactivation during NREM sleep, which are rapidly decaying across the first 40 minutes of Non-REM sleep in similar dynamics than dorsal, ventral or dHPC-BLA reactivation^5,12,15^. Although quantitatively similar, the reactivation of the joint assemblies generated during the aversive vs reward conditions are qualitatively different: the reactivation of the aversive joint assemblies reflect more accurately the preceding experience compared to the reactivation of the reward assemblies. This effect depends on the strength of recruitment of the aversive assemblies during wakefulness and originates in the increased recruitment of vHPC shock-responsive neurons into the coordinated dorsal and ventral SWRs that support the joint reactivation. Coordinated SWRs following the aversive condition are also enriched in dorsal place cell sequence replay. As such, they represent a privileged time-window for binding spatial dHPC and emotional vHPC offline representations with heightened accuracy. As in most previous studies tackling sleep-dependent processing of emotional experience in the dorsal hippocampus and other emotion-related structures^5,15^, we did not find any reactivation during REM sleep, despite evidence of its role in emotional processing^51,52^. It is possible that the methods used to assess reactivation are not appropriate to the intrinsic dynamics of REM sleep, or that the short epochs of REM sleep are drastically reducing the statistical power. Alternatively, REM sleep might contribute to emotional processing through other mechanisms than reactivation, such as synapse pruning and firing rate homeostasis^53,54^.

The salience of positive and negative stimuli cannot be perfectly matched, and the experimental design of the task to reach similar alternation behavior in the two conditions constrained us to use rewards at fixed locations and shocks at varying locations. We also recorded at plateau performance to maximize data collection. Thus, it is possible that the lack of increase in peak reactivation strength for the reward condition is due to the high familiarity of the animal with the task and maze, and would emerge following the exploration of a new environment. The increased accuracy we observed in the reactivation of the aversive assemblies may also be due to a higher salience of the shock, or to their less predictable nature. The vHPC was shown to be involved in both positive and negative emotional processing^35,36^, but its role in processing the negative end of the valence spectrum has been more firmly established^22,23,26,27^. It is therefore also possible that the increased accuracy reflects an inherent “fidelity” bias of the hippocampus towards negative information, reflected in the correlation between reactivation accuracy and recruitment strength during wake only for the aversive assemblies.

SWRs also occur in wakefulness during non-locomotor behaviors, such as pausing for reward consumption. In that case, dorsal and ventral SWRs differentially activate neurons in a common structure target, the NAc^35^, suggesting that independent dorsal and ventral SWRs can separately route spatial and positive emotional information to common efferent structures. During sleep without prior learning, a small proportion of SWRs, yet higher than during wakefulness, are coordinated between the dorsal and ventral poles^33,35^. We find that after an emotional spatial experience, this marginal subset of coordinated SWRs supports the reactivation of dorso-ventral neuronal assemblies. Because during coordinated SWRs the dSWR tends to precede the vSWR, we suggest that the extensive network of CA3 pyramidal cell collaterals and overlapping excitatory-inhibitory networks in CA1 or the direct projections from dCA2 to vCA1^55^ might allow for a subset of dSWRs to trigger vSWRs. A common input from the entorhinal cortex, subiculum or thalamic nuclei might also synchronize dorsal and ventral SWRs^2,3^.

The vHPC shows intricate bidirectional connectivity patterns with the emotional network^3^ that intersect with the functional diversity of vHPC cells^24,36^ as exemplified by the preferential connectivity of vHPC shock-responsive cells with the basal amygdala^26^. The synchronization of dHPC and vHPC SWRs, associated with heightened spatio-emotional precision, may account for previous reports of coordinated activity and replay between the dHPC and other regions, notably the basolateral amygdala and prefrontal cortex, despite the absence of direct anatomical connectivity^12,15,16,48^. Consistent with the literature^33,56^, coordinated dvSWRs show a higher amplitude than isolated dSWR or vSWRs, suggesting that they differentially affect the activity brainwide and in the direct output structures^57,58^.Thus, dHPC–vHPC coordination emerges as a potent mechanism for integrating and broadcasting spatial and emotional representations, consistent with the established role of SWRs in memory consolidation and their widespread effects on the rest of the brain^8^.

## Contributions

JFM and GG conceptualized the study, JFM, EP and ILP performed the experiments, JFM and AS analyzed the data. JFM and GG wrote the manuscript with contributions from all other authors.

## Acknowledgments

We thank Dr. Ralitsa Todorova, Dr. Nikolas Karalis, Katarína Studeničová, Olivier Peron and all Girardeau lab members for insightful discussions and advice on analysis. We are grateful to Laura Clément and Marie Sarraudy for assistance with pilot experiments, to Dr. Billel Khouader for his contribution to the experimental setup and to Baptiste Lecomte, Eloïse Marsan and all the zootechnician team for animal care.

## Material and Methods

### Ethical Statement

All experimental procedures were performed following the official European guidelines for the care and use of laboratory animals (86/609/EEC), the Policies of the French Committee of Ethics (Decrees n° 87-848 and n° 2001–464), and after approval by ethical committee (APAFIS #35991-2022031810445522 v3).

### Subjects and electrode implantation

This project included six individually housed female Long-Evans rats maintained on a 12h:12h light-dark cycle. Animals (300g/3 months-old at the time of the surgery) were injected with buprenorphine (Vetergesic® - 0.3mg/ml, dose: 0.05mg/kg) and deeply anesthetized with isoflurane. The local anesthetic Lidocaine (Lidor®- 20mg/mL, dose: 4mg/kg) was subcutaneously injected on the scalp 15 minutes before skin incision. The body temperature was monitored throughout the surgery and maintained at 37°C using a heating pad. Small metal screws were placed at different points in the skull to anchor the implant, and 2 screws were implanted above the cerebellum to serve as Ground and Reference. Two or three electrodes (NeuroNexus® Buzsaki64L for the vHPC, Buzsaki 32 or tetrode array for the dHPC) mounted on individual movable nano-drives (Cambridge nano-drives, size: 2×5×11mm) were implanted above the vHPC unilaterally (initial position, AP: -5.50mm, LAT: 4.10mm, DV: -6.5mm, angle: 10°) and in the dHPC either unilaterally or bilaterally (initial position, AP: -3.50mm, LAT: 2.50mm, DV: -1.7mm). The drives were secured on the skull using dental cement. The drives and probes were protected by a cement-covered copper-mesh Faraday cage on which the probe connectors were attached. For the eyelid stimulation, a bipolar electrode made of two 140 μm silver wires was inserted between the skin and the muscle surrounding the eyelid. The connectors were fixed to the copper mesh. Following the surgery, animals were injected with a combination of Buprenorphine (Vetergesic® - 0.3mg/ml, dose: 0.05mg/kg), and Meloxicam (Metacam®-5mg/mL, dose: 1mg/kg) at 24 and 48h. Animals were allowed to recover for at least 5 days with *ad libitum* food and water before training started. In one animal, the dHPC electrode failed to reach the pyramidal layer and only vHPC data was used.

### Recordings and Behavior

All data will be made available upon reasonable request to the first or corresponding author. All animals were free from prior manipulation before being included in the study. All experiments were performed during the light cycle. After at least 5 days of daily handling, animals were water-deprived and trained to run back and forth on a linear track for water rewards. After 7 days of training, animals regained access to *ad libitum* water and underwent electrode implantation surgery 3 days later. After the recovery period, animals were water-deprived again and re-exposed to the rewarded linear track task for at least 2 days. Subsequently, animals were exposed to three daily training sessions composed of a rewarded and an aversive linear track exploration. For pre-training on the aversive run condition, animals were initially provided water at the end of each lap for a certain time, while the remainder of the time, they received an electric shock to the eyelid if they remained motionless for at least 20 seconds. The reward time was progressively reduced across training days until only shocks were delivered if the animal remained quiet. Once the animals were pre-trained, task sessions were composed of pure reward and pure aversive run sessions, preceded and followed by sleep sessions (**Fig 1a-b**). Across days, the Aversive and Reward runs were presented in a pseudo-randomized order, ensuring that both the Aversive-Reward and Reward-Aversive combinations occurred approximately the same number of times over the course of the experiment. At the end of each recording day, animals had access to *ad libitum* water for 5 to 20 minutes to ensure the animals remained at >80% of their normal weight. During pre-training, the probes were slowly lowered in the brain to reach the dorsal and ventral CA1 pyramidal layers. Afterward, the probes were adjusted daily to optimize ripple and unit recording. Signals were recorded at 20 kHz using an Intan recording system (Intan Technologies, California, United States of America) and videos captured at 30 fps with a Basler camera (ace Basler 106754) synchronized with the Intan system. The instantaneous speed was calculated using the neck position extracted from the videos using DeepLab Cut (DeepLabCut2.0 Toolbox)^59^.

### Preprocessing

Electrophysiological data was sampled at 20 kHz and down-sampled to 1250 Hz to extract the local field potentials (LFPs). Spikes were extracted by high-pass filtering and thresholding the signal, then clustered using Kilosort 2.5 followed by manual curation using Phy. Sleep stages were manually scored through visual inspection of the dorsal and ventral hippocampal spectrograms and accelerometer signals using TheStateEditor, a custom program from the FMA toolbox (http://fmatoolbox.sourceforge.net/). We defined NREM epochs as periods characterized by immobility and low theta/delta ratio, while REM sleep was characterized by immobility and regular theta waves (**Extended data Fig. 11a-c**). Ripple detection was performed by band-pass filtering (∼100–200 Hz), squaring, z-scoring, and thresholding the LFP recorded in the dorsal or ventral CA1 pyramidal layer. Ripples were defined as events lasting between 20 and 100ms, starting at 2 or 3 standard deviations (s.d.) above the mean, and peak amplitude exceeding 5 or 7 s.d. for dorsal or ventral ripples, respectively.

### Statistics and Analysis

All analyses were performed using customized toolboxes (Chronux, FMA toolbox), built-in functions, and custom-written scripts in Matlab (The MathWorks, Inc., Natick, MA, USA). All codes used to generate the figures are available on the Girardeau Lab github repository (https://github.com/GirardeauLab/Morici_et_al). If the data met the criteria for normality, we used Student’s t-tests for two-group comparisons and one-way ANOVA followed by the Tukey-Kramer post-hoc test for multiple-group comparisons. A two-way ANOVA was used to assess the effects of two independent variables, each with multiple levels, followed by Bonferroni-adjusted Tukey-Kramer post-hoc tests for pairwise comparisons between groups. If data did not meet normality criteria, we used a two-tailed unpaired Mann-Whitney test for comparisons between two groups. For more than two groups, we used the Kruskal-Wallis test for unpaired data and the Friedman test for paired data, followed by the Tukey-Kramer post-hoc test for pairwise comparisons between groups. Data presented in cumulative graphs were compared using the Kolmogórov-Smirnov test. Correlations between continuous variables were examined by linear regression. A value of p<0.05 was considered statistically significant unless otherwise stated. Data were presented as boxplots, where the edges of the box represent the 25th and 75th percentiles and the central line indicates the median, or as scatterplots showing individual data points together with the mean.

### Dorsal-ventral ripples coordination

First, we constructed a cross-correlogram of ventral SWR peaks locked to dorsal ones using the *CCG* function from FMAtoolbox and normalized it to the total amount of ventral SWRs (10-msec bins). By visually inspecting the cross-correlogram, we defined coordinated events as dorsal-ventral ripple pairs occurring +/-100ms from each other. To ascertain whether the percentage of coordinated dorsal and ventral ripples exceeded the chance level, we shuffled the reference ripple peaks 100 times and calculated the percentage of coordinated reference ripples. This process yielded a surrogate average percentage value compared with the value obtained in each session. For auto-correlograms and cross correlograms in Figure 6, we used 10-msec bins. Directionality inside each dorsal-ventral coordinated ripples pair was calculated as the percentage of coordinated ripple events in which the dorsal ripple preceded the ventral one and vice-versa (**Extended Data Figure 17**).

### Single-units analyses

We detected 1071 single units in the dHPC and 544 in the vHPC in 6 animals. Units were classified into putative pyramidal cells and putative interneurons based on waveform shape (**Extended data Fig. 4**): duration at 50% from baseline to peak (distance A) and peak-to-trough time (distance B) were used for clustering with k-means into 2 clusters, interneurons, and pyramidal cells. The gain locked to ripple peaks was calculated in a 10-msec bin by dividing the firing rate in each bin by the baseline firing rate outside SPW-Rs (inter-ripple NREM intervals). To define if a unit was ripple-modulated, we performed a Poisson test with P < 0.001 (**Extended data Fig. 11d-e**). For **Fig. 2C**, Principal Component Analysis (PCA) was performed on the spike trains of all dorsal and ventral hippocampal neurons separately. Since the number of PCA dimensions is related to the number of neurons recorded, we only included sessions with at least three single units. We concatenated periods when the animal was exploring the environment (>7 cm/sec for more than 2 seconds) during aversive and reward run sessions. We then measured the Euclidean distance between the centroids of the aversive and reward projections^60^.

Place cells were defined in the dHPC and vHPC as putative pyramidal cells with spatial information content^61^ exceeding the 90th quantile of the chance distribution and the presence of at least one place field in at least one of the two run conditions. Chance distribution was assessed by circularly permuting position data relative to spike times for each unit. To construct the place tuning curve, position data were discretized into 2.5 cm bins, and rate maps were computed as smoothed firing-rate histograms (Gaussian kernel, sigma = 2) normalized by occupancy time per bin. Data from the ends of the linear track, where rewards were delivered, were excluded from the analysis. Only periods during which the animals maintained speed higher than 2.5 cm/s were considered. Place fields were defined as contiguous regions comprising a minimum of four spatial bins (10 cm) that contained the location of the maximum firing-rate peak (≥1 Hz). Within these regions, firing rates were required to exceed 60% of the peak firing rate value.

To evaluate remapping, we calculated spatial correlations as bin-by-bin Pearson’s correlations of the rate maps, both within and between conditions. For within-condition comparisons, position and spike data from a given condition were divided into *n* bins of 1 s each, and bins were randomly assigned into two groups of size *n*/2. Separate rate maps were generated for each group, and their spatial correlation was computed. This procedure was repeated 100 times per cell, and the mean correlation was reported. For between-condition comparisons, data from each condition were similarly divided into two groups to construct a rate map. To compute the spatial correlation one of the two rate maps from each condition was randomly chosen. This process was repeated 100 times per cell, and the mean was reported^62^.

To assess the unit response to shocks or rewards, the firing rate of each single unit was computed using 100-ms time bins in a (±5 seconds) centered around the time of shock delivery or reward delivery and z-scored. The reward was delivered upon the opening of the water valve triggered automatically by the passage of the rat through infrared detectors placed at the entrance of the track end platform (see schematic Fig. 3h). The average delay from reward delivery to reaching reward location was 0.93±0.02 sec. To identify reward or shock-responsive units, a surrogate distribution was generated by averaging the responses following the shock/reward after shuffling the spike times 200 times. Units with a firing rate exceeding the 90th percentile of this surrogate distribution were classified as shock- or reward-responsive. To identify speed-modulated units, linear correlations between firing rates and mean speed outside of the shock periods were calculated in 1-second bins. Units with a p-value < 0.05 were considered speed-modulated. To remove the linear influence of speed on neural responses during shock delivery or approaching the reward port, the Gram-Schmidt orthogonalization process was applied^63,64^. Briefly, the mean neuronal activity was projected onto the mean speed trace, and the residual component was used for subsequent analysis. For statistical comparisons and plotting (Fig. 3a,d,e,f,e,j) of the response during the shock or reward before and after speed decorrelation, the maximum response value during the shock was used after each trace was normalized to its baseline period (from -5 sec to -1 sec before the shock or reward).

### Response during coordinated ripples

The firing rate of each pyramidal cell was calculated in 10-ms bins and smoothed using a 2SD Gaussian kernel. Each tuning curve was then normalized by the mean firing rate during NREM sleep outside of ripple events. For coordinated and uncoordinated ripples, the peri-event firing rate was calculated within ±500 ms of the ripple event. These firing rate curves were further normalized by the mean firing rate of each tuning curve, calculated in a time window ±200 ms from the event. The firing activity from dorsal and ventral pyramidal cells was locked to the coordinated dorsal or ventral ripple member, respectively. Statistical comparisons of pre- and post-NREM ripple mean gains were performed using a one-sided Mann-Whitney test on the maximal gain in a ±300-ms window centered at the ripple peak. To correlate the shock-induced response with the response during sleep, the difference between the average response during coordinated ripples in post-NREM and pre-NREM sleep was linearly correlated with the maximal response during shock delivery in the aversive run.

### Assembly Analysis

Assembly detection was performed only in sessions having at least 3 neurons in the dorsal or ventral hippocampus that were active in both conditions. We binned and z-scored the spike trains of dorsal and ventral pyramidal cells during mobility periods (>7 cm/sec during more than 2 seconds) into 25ms bins, considered the optimal bin for assembly detection^7^. Assembly patterns were identified using a two-step process using ICA and PCA^43^ (**Extended data Fig. 7**). After detecting the assembly patterns, units with weights exceeding the threshold defined as abs(1/√N) were defined as assembly members. Assemblies with members in both the dHPC and vHPC were defined as joint assemblies.

### Assembly patterns analyses

To track assembly pattern expression over time (**Extended data Fig. 7**), a projection matrix was created for each pattern using its weight vector’s outer product, with the main diagonal set to zero. To maximize the role of members in assembly activations, we set to zero the weight of non-member units. Each neuron’s spike train was convolved with a Gaussian kernel and z-scored. The expression strength of a pattern at any time was defined by the quadratic form of its projection matrix with the smoothed and z-scored firing rate vector. A peak was defined as strength>5. We calculated the peak rate in 60-second windows during Pre- and Post-sleep. To assess assembly activation dynamics in the real-time course of post-NREM sleep, peak rates were normalized to the average rate during pre-NREM sleep and aligned to the onset of the first post-NREM epoch. To minimize the overlap between the Post-sleep of the first emotional condition and the Pre-sleep of the second one, we restricted NREM or REM epochs to those occurring 1 hour before or after each run session. Peak Reactivation Strength was calculated as the average values of peaks occurring during Post-sleep NREM minus the average of the peak values in Pre-sleep NREM divided by the average value of all peaks detected during both sleep sessions. The calculation was restricted to peaks occurring during NREM periods or Ripples.

### Replay detection

The replay sequence detection algorithm was implemented using the code provided by Tingley & Peyrache (2020)^65^. Candidate replay events were defined as dorsal ripples containing at least three active Place-cells. To evaluate whether these events carried spatial information, we first applied a Bayesian reconstruction method. Briefly, given the average firing rate *f _i_*(*x*) for n dorsal hippocampal neurons, we estimated the animal’s location *x* given the number of spikes *n* of all the cell within a time window *τ*.

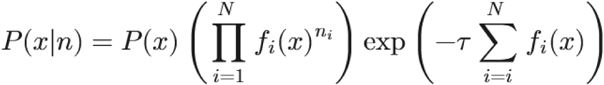

where n is the number of dorsal hippocampal place cells and P(x) is the probability of a certain position being reactivated. As a complementary approach, we used a previously described template-based rank-order correlation method^46,65^. This method correlates the order of place-field activation on the linear track with the spike sequence observed during a candidate replay event. Candidate replay events were discretized into 20-ms bins across the ripple duration, producing a posterior probability matrix (position × time). To identify sequential trajectories, the Radon transform^66^ was applied to this matrix, and the integral of the best-fit line was calculated to quantify the strength of linear trajectories consistent with replay. For both methods, significance was determined by comparing observed values to surrogate distributions obtained by circularly shuffling place-cell ids. A candidate was classified as significant if it exceeded the 90th percentile of its corresponding surrogate distribution. Because uncoordinated ripple events were more frequent than coordinated ones, they were subsampled 1000 times, and the mean percentage of significant replays was computed.

## Extended Data

**Extended Data Figure 1:**
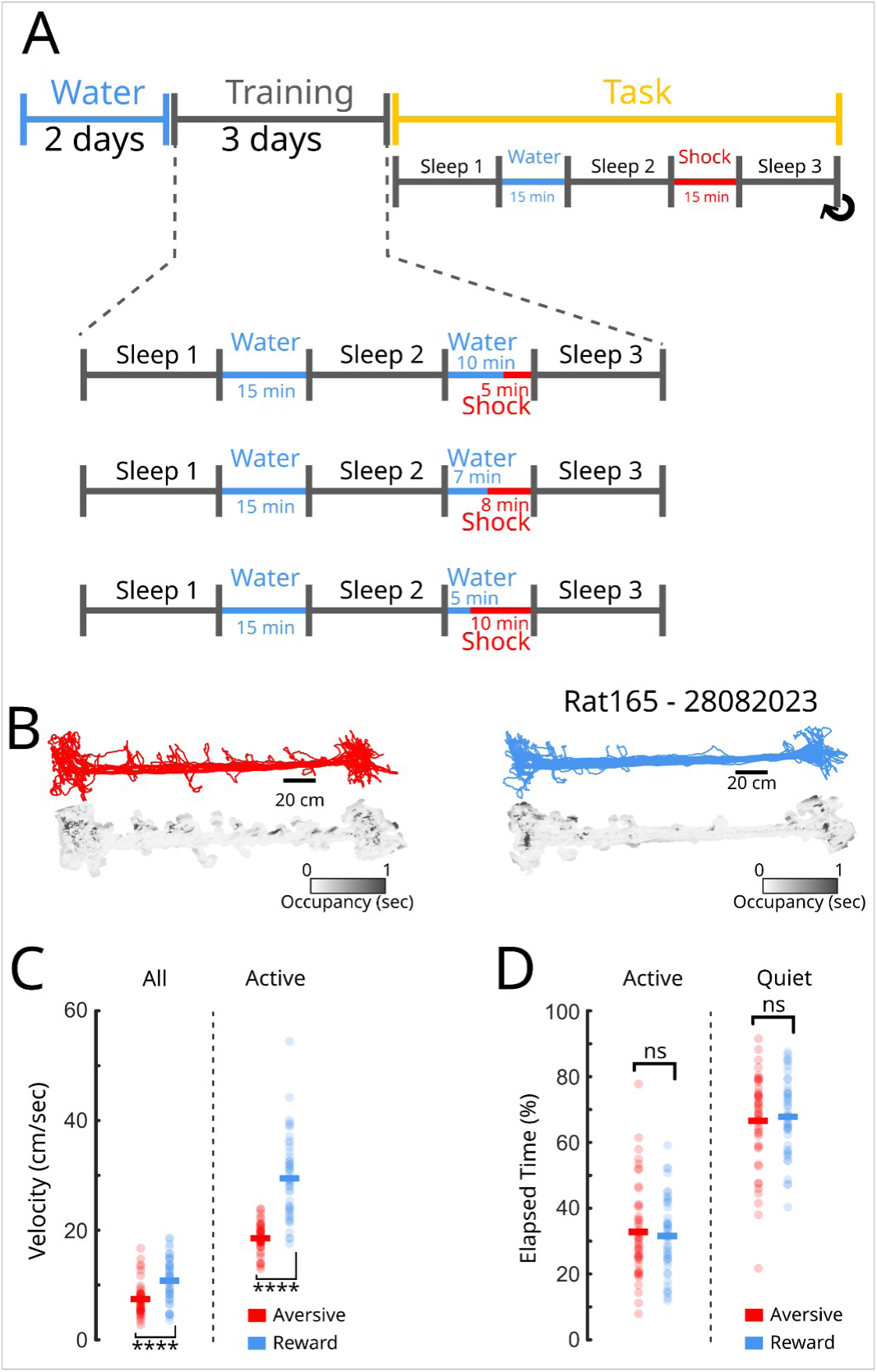
Training protocol and behavioral characterization. **A**, After the implantation surgery, animals were re-trained to perform the rewarded run (“Water”). Then for 3 days, the shock was gradually introduced and water rewards eliminated to train the animals on the aversive run (“Training”). For the following days, animals performed the rewarded and aversive runs on the same day, starting with the rewarded or aversive run in alternation (Task). The recordings started when they reached plateau performance for both run types (see Main text and Fig.1). **B**, Examples trajectories and occupancy maps (grey) for aversive (red) and rewarded-motivated exploration (/blue). Active periods were defined as moments where the animals showed an instantaneous speed of >7cm/sec for at least 2 seconds. **C**, Average velocity for aversive and rewarded-run sessions during the whole run (left) or during the active period only (right). (two-sided t test, ****<0.0001). **D**, Time spent in the active and quiet state on the track. (two-sided t test, p=0.62, N_sessions_=47).

**Extended Data Figure 2:**
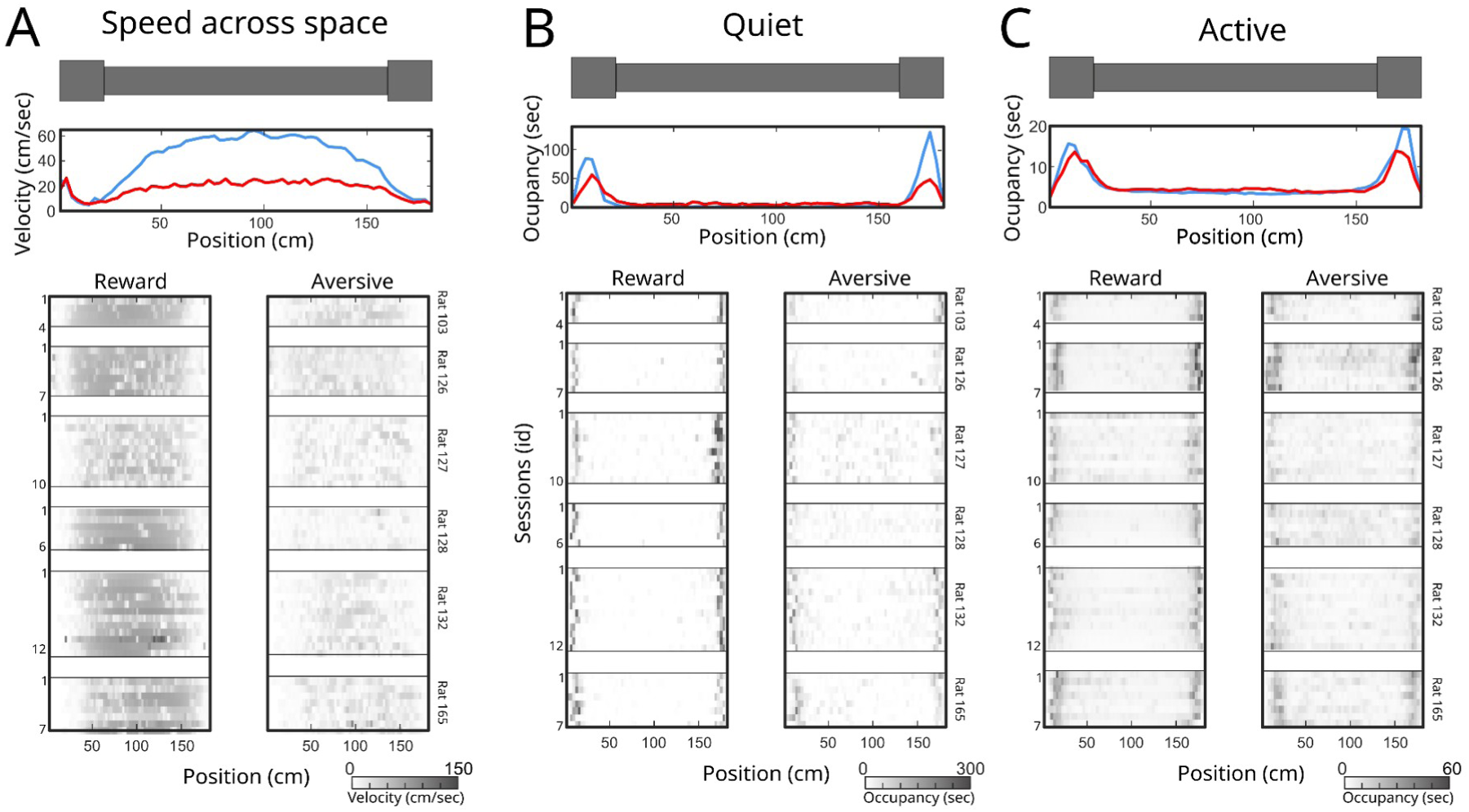
Distribution of speed, quiet, and active periods across the linear track. **A**, Average speed as a function of location on the track for all aversive(red; n=47) and rewarded runs (blue; n=47; top) and for individual runs (greyscale; bottom). **B**, Average time spent in quiet behavior (<7cm/sec) as a function of location on the track for all runs (top) and individual runs (bottom;greyscale). **C**, Same as **B** but for active behavior.

**Extended Data Figure 3:**
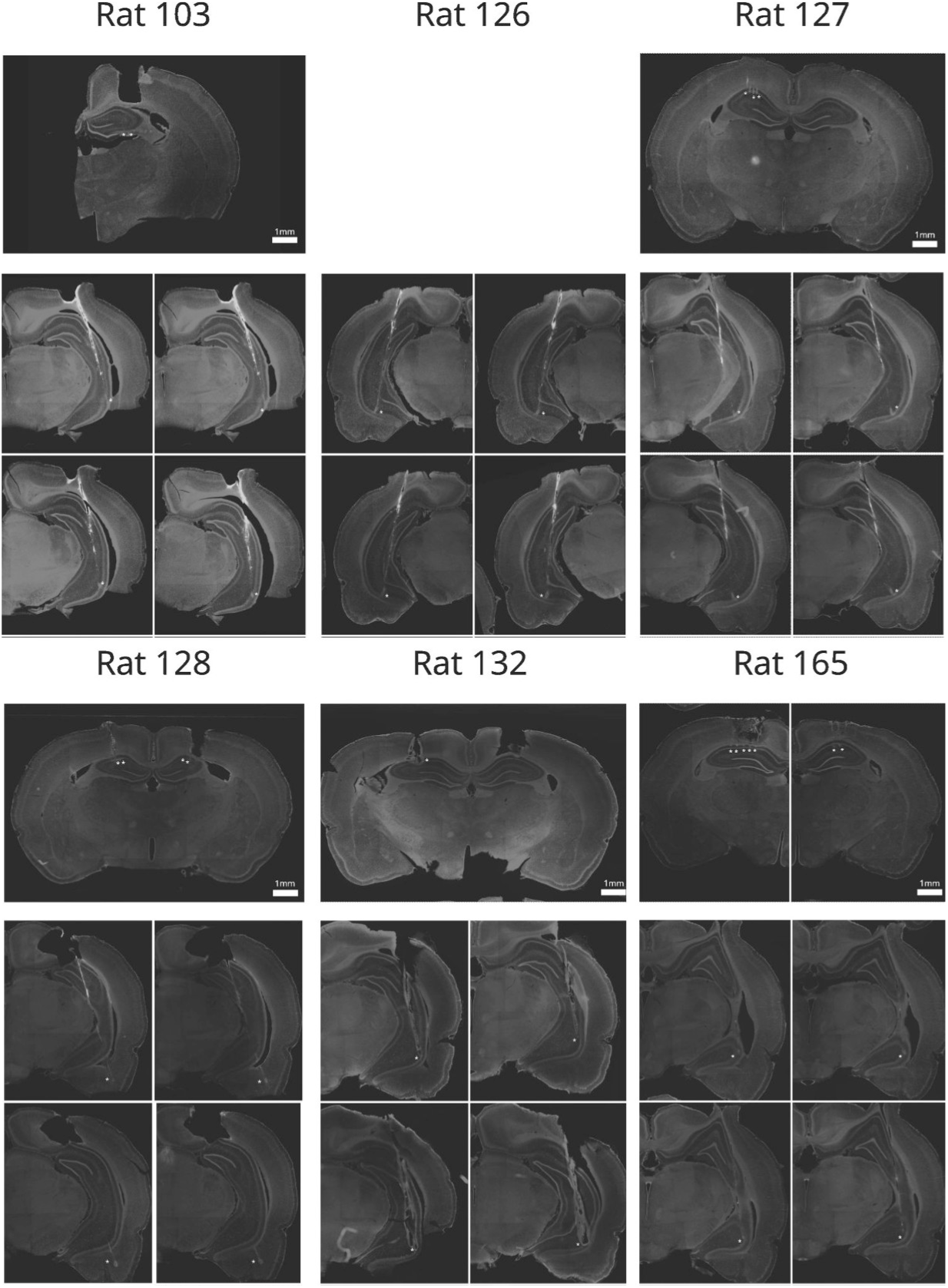
Histological reconstruction of successful electrode placement in dorsal and ventral hippocampus. In five rats (Rats 103, 127, 127, 132, 165), we reached both the dorsal and ventral pyramidal layer. In one rat (Rat 126), the dorsal electrode did not reach the pyramidal layer.

**Extended Data Figure 4:**
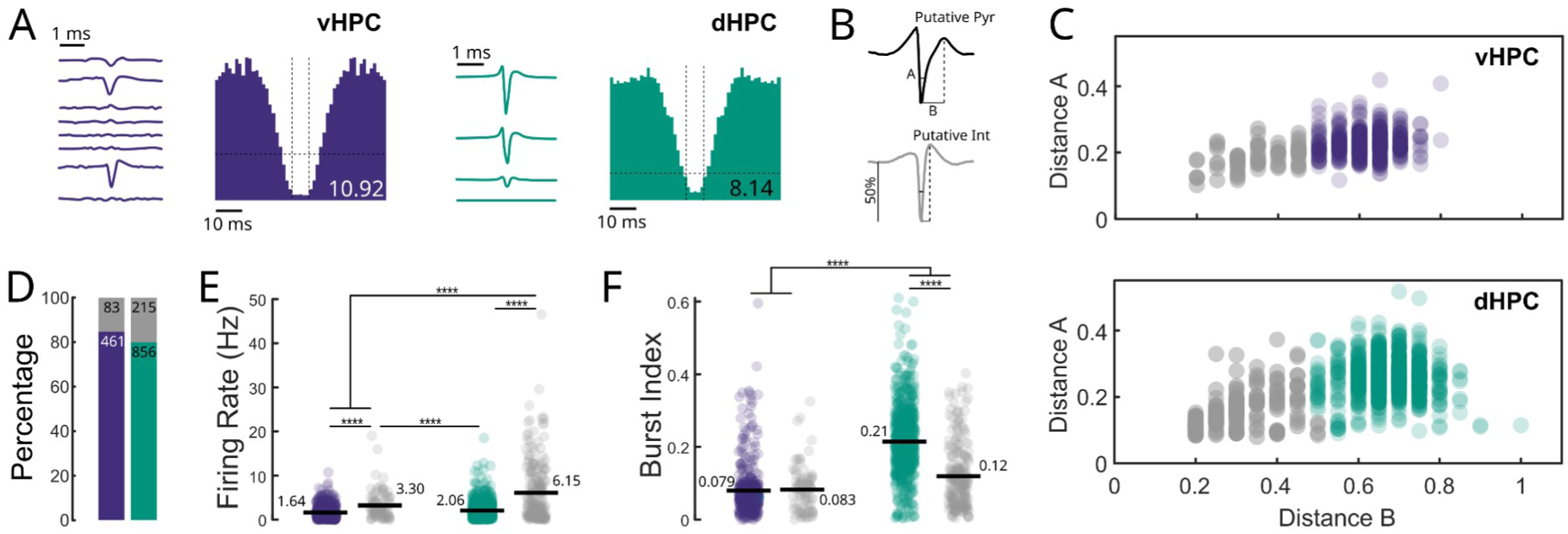
Single-units waveform classification and characterization. **A**, Examples of ventral (left panel, purple) and dorsal (right panel, green) single units. **B**, Units were classified into putative pyramidal cells (Pyr) or putative interneurons (Int) using k-means clustering with 2 clusters on duration at 50% from baseline to peak (distance A) and peak-to-trough time (distance B). **C**, The clustering into putative pyramidal cells (green/purple) and putative interneurons (grey) was performed separately in the vHPC (top) and dHPC (bottom). **D,** Percentage of Pyr (green/purple) and Int. (grey) in the dHPC (green, N_int_:215, N_pyr_=856) and vHPC (purple, N_int_:83, N_pyr_:461). **E**, Firing rates of putative Pyr and Int. in the vHPC (purple) and dHPC (green). (Two-way ANOVA, p_cell_type_<0.0001, F=177.73, p_structure_<0.0001, F=57.73, p_interaction_<0.0001, F=31.76, Bonferroni post-hoc test, ****p<0.0001). **F**, Burst index for Pyr. and Int. in the vHPC and dHPC. (Two-way ANOVA, p_cell_type_<0.0001, F=177.73, p_structure_<0.0001, F=57.73, p_interaction_<0.0001, F=31.76, Bonferroni post-hoc test, ****p<0.0001).

**Extended Data Figure 5:**
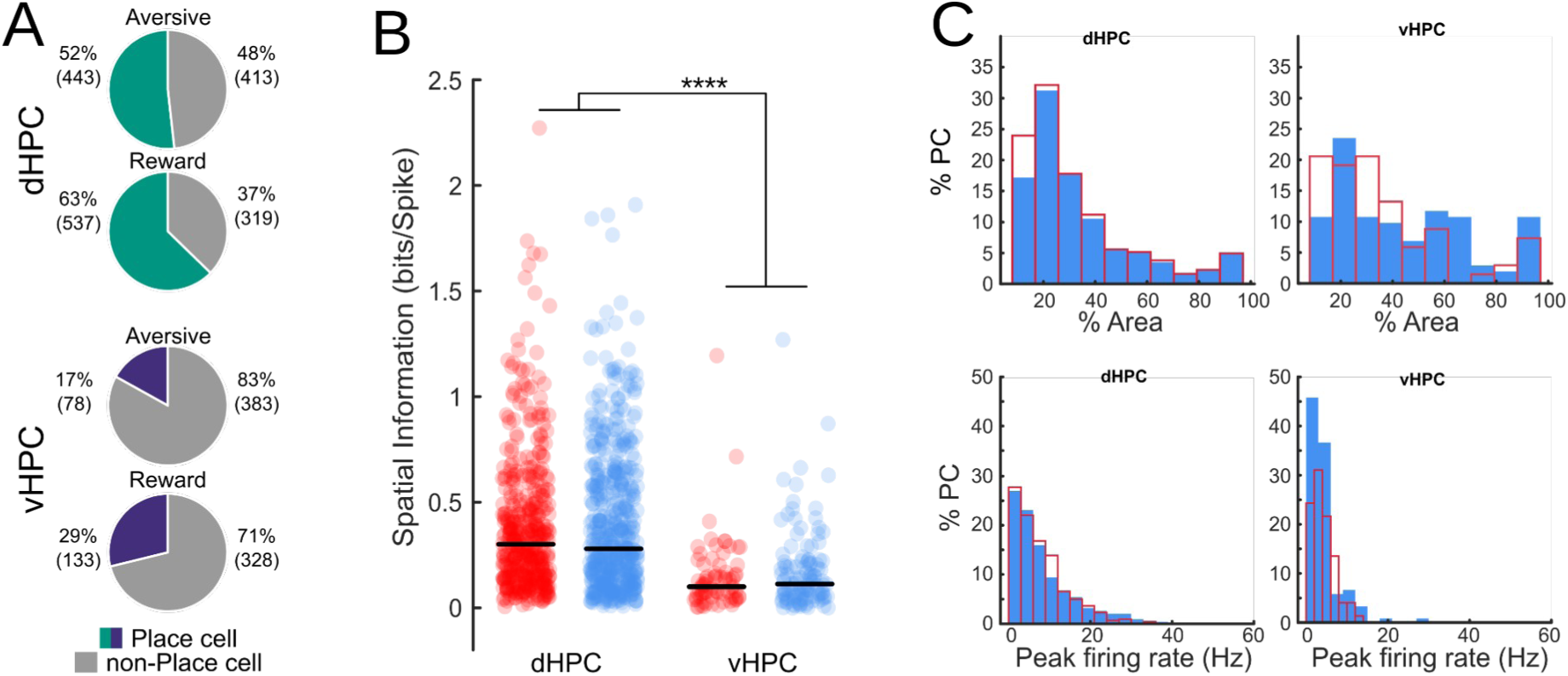
Spatial Properties of dorsal and ventral place cells across conditions. **A**, Proportion of place cells among dorsal and ventral pyramidal neurons across emotional conditions. (Chi^2^, dHPC: ***p=0.00001, vHPC: ***p=0.00002). **B**, Spatial Information for dorsal (left) and ventral (right) place cells during aversive (red) and reward (blue) sessions (Two-way ANOVA. p_interaction_=0.71, p_condition_=0.48,****p_structure_<0.0001). **C**, Place cell characteristics across the dorsal-ventral hippocampal axis. Upper panel, Percentage of the area of the maze covered by the place-field in dHPC (left) and vHPC (right). Solid-bars histogram represent place-cells from reward run sessions, while empty-bar histogram represent those ones detected during aversive run sessions (two-sample Kolmogorov-Smirnov test, dHPC: p=0.06, vHPC: p=0.06). Lower panel, Peak firing rate for dorsal (right) and ventral (left) place-cells detected during the rewarded (blue) or aversive (red) condition (two-sample Kolmogorov-Smirnov test, dHPC: p=0.22, vHPC: p=0.35).

**Extended Data Figure 6:**
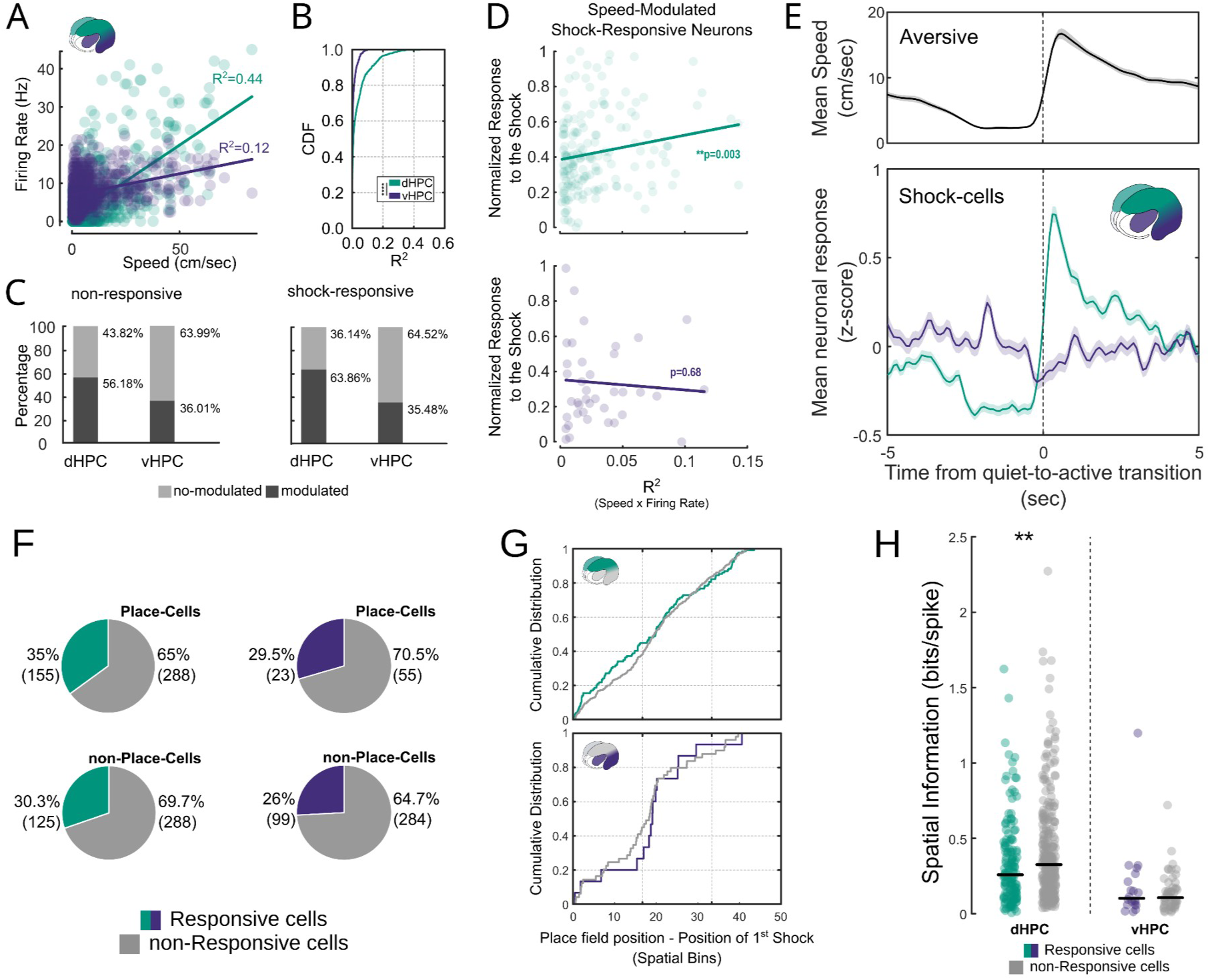
Speed, shock and place-modulation of activity in aversive run. **A**, Linear correlation between the firing rate of 2 example putative pyramidal neurons and the animal’s speed in the dHPC (green) and in the vHPC (purpleblue). **B**, Cumulative Density Distribution of R-Square Linear Correlations for Dorsal and Ventral Hippocampal Neurons. Kolmogórov-Smirnov test,, ****p<0.0001. **C**, Percentage of speed-correlated and uncorrelated activity of shock-responsive and non-responsive neurons. (non-responsive, Chi^2^, **p=0.003; shock-responsive, Ch^2^, ****p=0.00004). **D**, Linear correlation between the R-squared values of Speed-Neuronal activity in shock-responsive neurons. (Upper panel: dHPC, R^2^=0.052, p=0.003, n=170; Lower panel: vHPC, R^2^=0.004, p=0.68, n=40). To ensure that the observed correlation in the dHPC was not due to the larger number of neurons, we subsampled the dHPC population 1,000 times to match the size of the vHPC population and performed linear regression. The R^2^ value obtained from the entire population was not significantly different from the distribution generated by subsampling (Kolmogórov-Smirnov test, p=0.67). **E**, Activity of dorsal and ventral shock-responsive neurons during quiet-to-active transitions. Upper panel, Average speed across all transitions of aversive sessions(N_sessions_=47, N_transitions_=45-108). Lower panel, Normalized firing of dorsal (green) and ventral (purple) shock-responsive neurons. Dorsal responses followed speed dynamics, whereas ventral responses were not modulated by locomotion. (N_dorsal_=285, N_ventral_=124). **F**, Percentage of shock-responsive (green: dHPC; purple: vHPC) and non-responsive (gray) cells, separated into place-cells and non–place-cells. (Chi^2^ test, Place vs. Non-Place: dHPC, χ^2^=2.16, p=0.14; vHPC, χ^2^=0.44, p=0.51). **G**, Cumulative distribution of the distance between the place-field position of shock-responsive place-cells (green/purple) and pure place-cells (gray). (Kolmogorov–Smirnov test , p_dHPC_=0.34 , p_vHPC_=0.76). **H**, Spatial information of shock-responsive vs. non-responsive place-cells. (Wilcoxon test, **p_dHPC_=0.008, p_vHPC_=0.67).

**Extended Data Figure 7:**
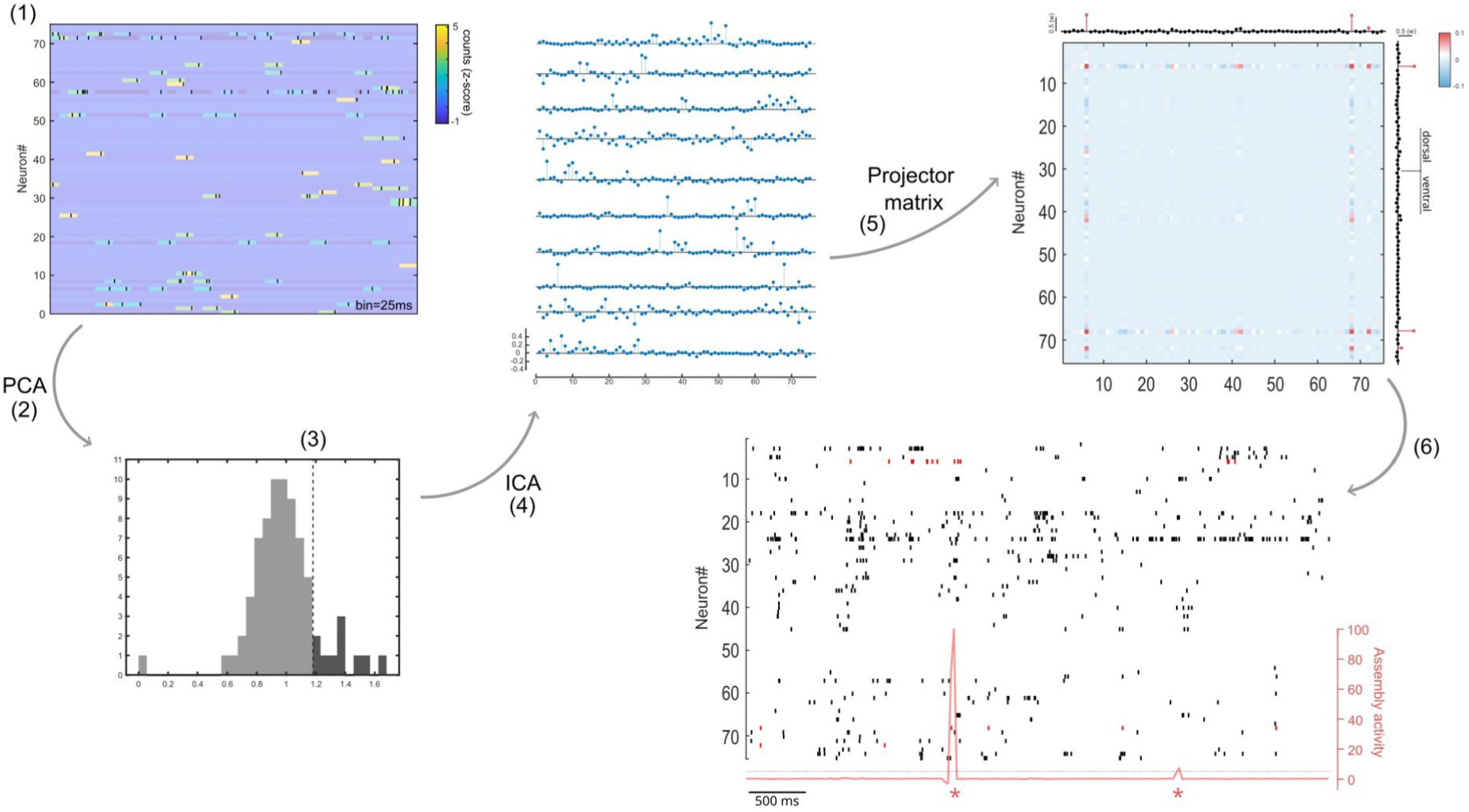
Schematic of the procedure for assembly pattern identification. . The procedure was performed using putative-pyr units coming from both the dorsal and ventral hippocampus. After z-scoring the spike trains (1, 25ms time bins) , a principal component analysis (PCA, 2) was performed. To identify the patterns associated with significant co-activations between units, the Marcenko-Pastur threshold was applied (3). Independent component analysis (ICA, 4) was subsequently applied to identify assembly patterns that were above the threshold applied in step 3. To track the assembly pattern activation, a projector matrix was constructed by calculating the outer product of the weight vector and setting the matrix diagonal to zero (5). The assembly activation strength was calculated as the quadratic form of the projector matrix with the convolved and z-scored spike trains (6).

**Extended Data Figure 8:**
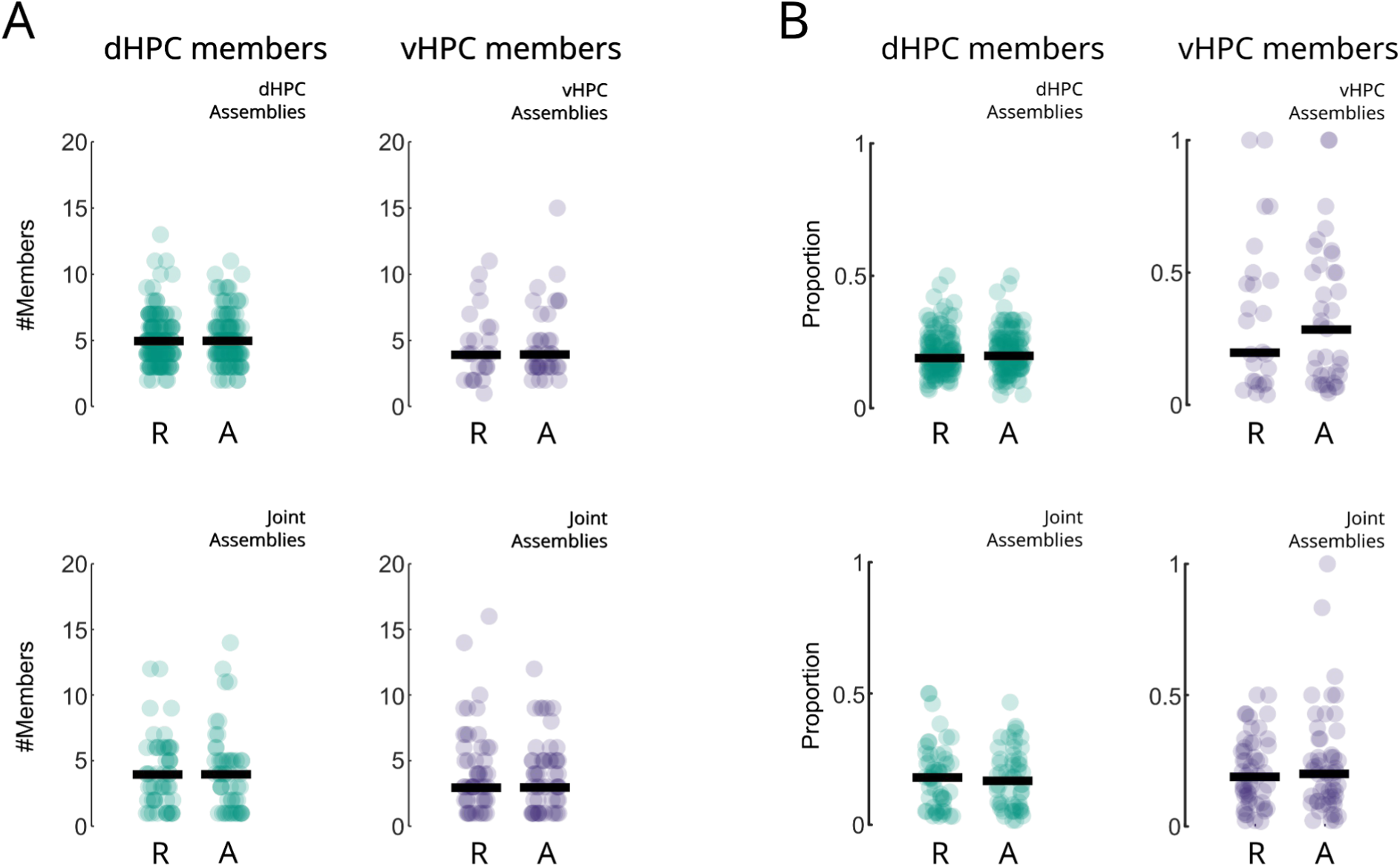
Number of significant members of the assemblies. Assembly members were defined as units for which the ICA weight exceeded a threshold defined as the absolute value of 1/√N_units_. **A**, Number of dorsal (left) and ventral (rigth) members in dorsal, ventral (top), and joint assemblies (bottom) detected during rewarded (R) or aversive (A) runs . (unpaired Wilcoxon test, dHPC Assemblies, p=0.55; vHPC assemblies, p=0.88, Joint Assemblies, p_dHPC_=0.79, p_vHPC_=0.70). **B**, Proportion of members relative to the total number of recorded units during the session. No significant difference was observed between aversive and reward assemblies.(unpaired Wilcoxon test, dHPC Assemblies, p=0.32; vHPC assemblies, p=0.96, Joint Assemblies, p_dHPC_=0.94, p_vHPC_=0.78). (two-sided Mann-Whitney test).

**Extended Data Figure 9:**
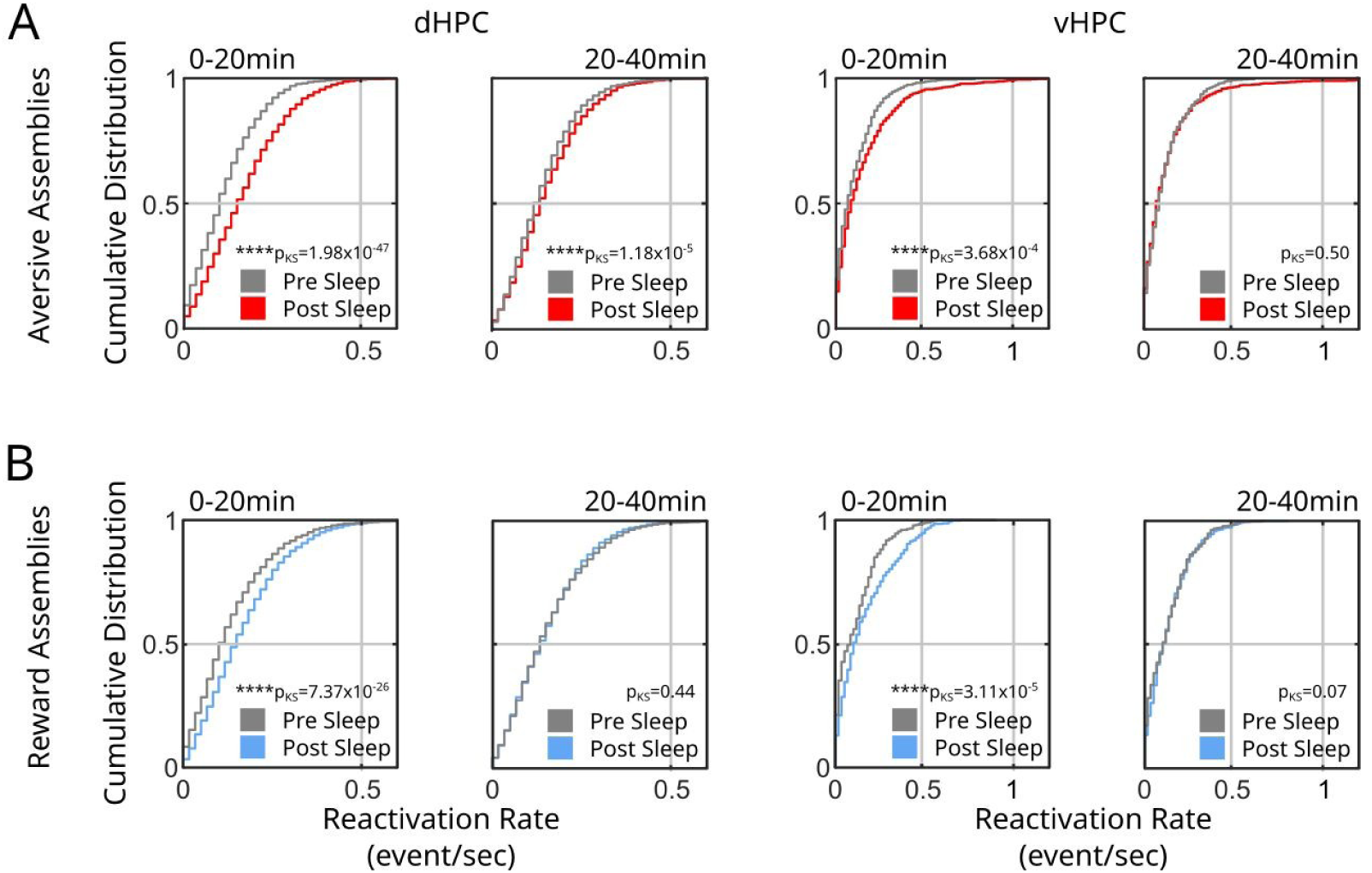
Reactivation rate of dorsal and ventral assemblies. Cumulative distribution of assembly reactivation rate in 60s bins during pre- and post-sleep sessions. The reactivation rate was calculated in 60-second bins aversive (**A**) and rewarded (**B**) assemblies’ activity. In both: dHPC and vHPC assemblies, the reactivation rate was significantly higher at the beginning of the post-sleep session and progressively went back to pre-sleep levels at later time bins. (Kolmogórov-Smirnov test, ****p<0.0001). Aversive assemblies, N_dHPC_=131, N_vHPC_=37; Reward assemblies, N_dHPC_ =146, N_vHPC_=25.

**Extended Data Figure 10:**
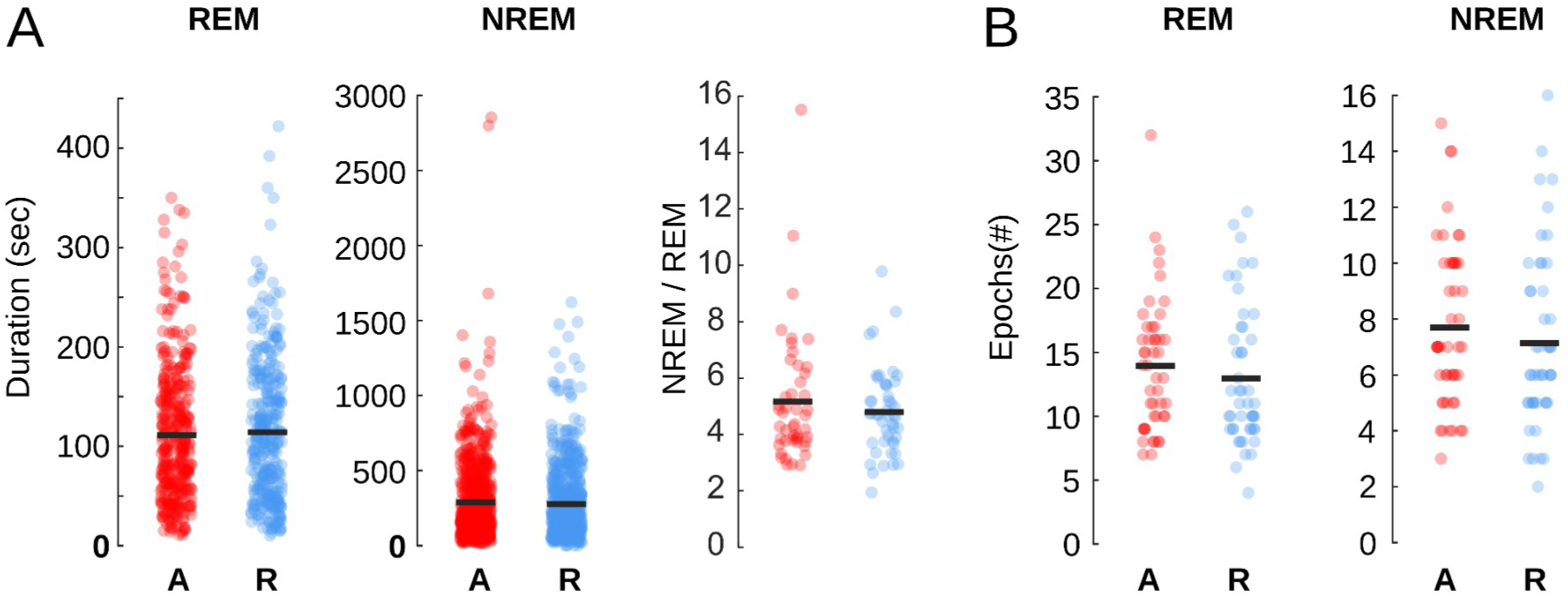
Emotional conditions do not influence general parameters of sleep architecture. **A**, REM and NREM sleep bout duration following aversive (red) and rewarded (blue) run sessions (unpaired Wilcoxon test, REM: p=0.64, N_aversive_=362, N_reward_=336; NREM: p=0.68, N_aversive_=655, N_reward_=611). Right panel, NREM/REM Ratio from all recorded sessions (unpaired Wilcoxon test, p=0.8’, N=47). **B**, Number of REM and NREM bouts following the aversive or rewarded contition (unpaired Wilcoxon test, REM: p=0.30; NREM: p=0.23, N_sessions_=47).

**Extended Data Figure 11:**
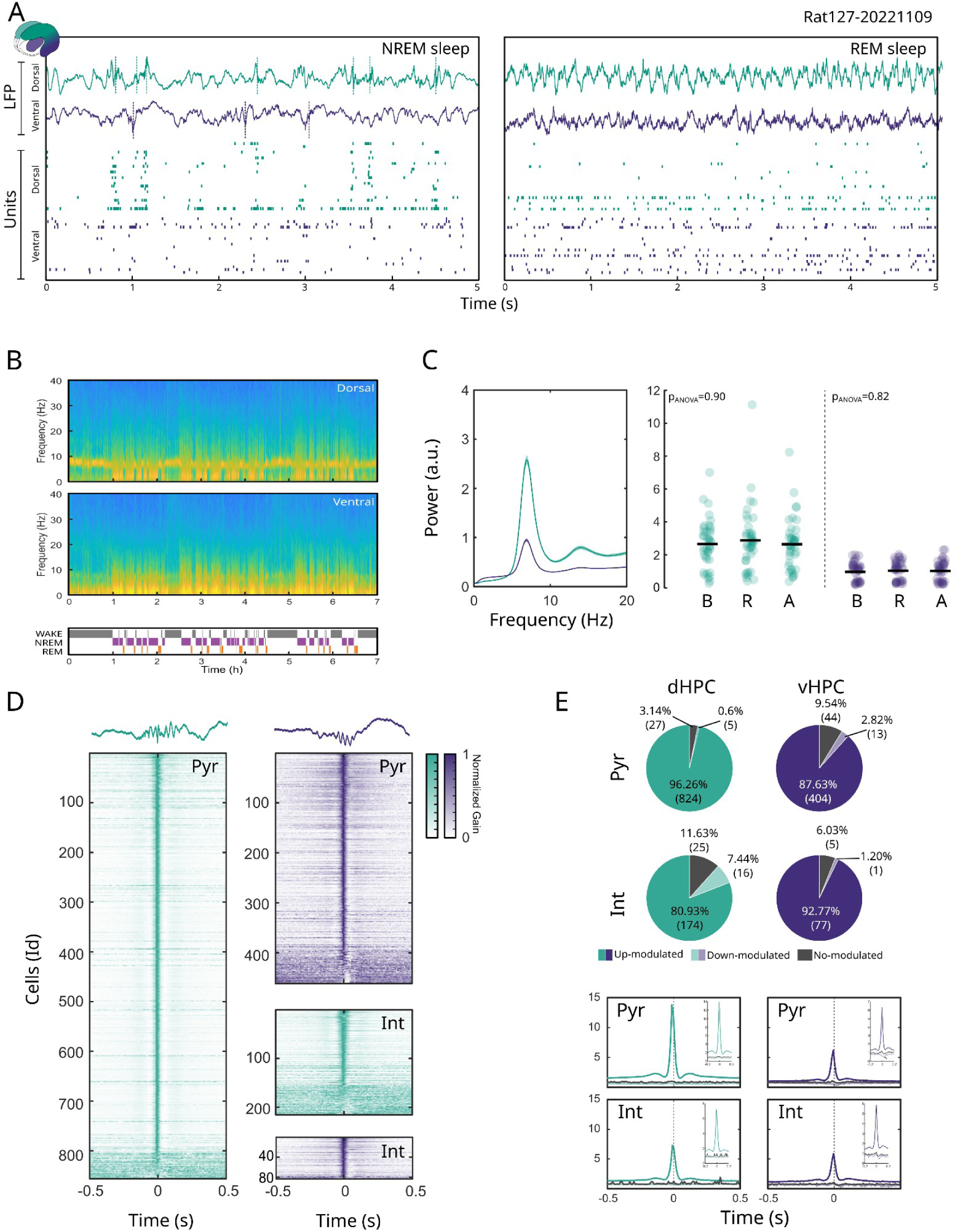
Sleep characterization in dorsal and ventral hippocampus. **A**, Representative example LFP and raster plots of single unit activity in the dHPC (green) and vHPC (purple) during NREM and REM sleep. Note the theta oscillation of the LPF during REM sleep and ripples (dashed line) during NREM sleep. **B**, Example spectrograms for a full session calculated on the dHPC (top) and vHPC (bottom) showing the changes in the various frequency bands depending on the corresponding state. **C**, Power spectrogram calculated from dorsal (green) and ventral (purple) hippocampal LFPs during REM sleep (left). Power spectrum maximum value detected in theta band during REM sleep (6-10 Hz) in baseline (B), post-rewarded run (R), and post-aversive run (A) sleep sessions. (One-way ANOVA, dHPC: p=0.90, N_dHPC_=40; vHPC: p=0.82, N_vHPC_=47). **D**, Normalized peri-event gain of dorsal (green) and ventral (purple) pyramidal cells and interneurons to local ripple events. E, Percentage of cells significantly up- or down-modulated or not modulated during local ripples (Poisson test, with p<0,001) for pyramidal cells and interneurons in the dHPC and vHPC (top). Average gain in the dHPC and vHPC for Pyr. and Int. grouped according to their modulation during ripples.

**Extended data Fig. 12:**
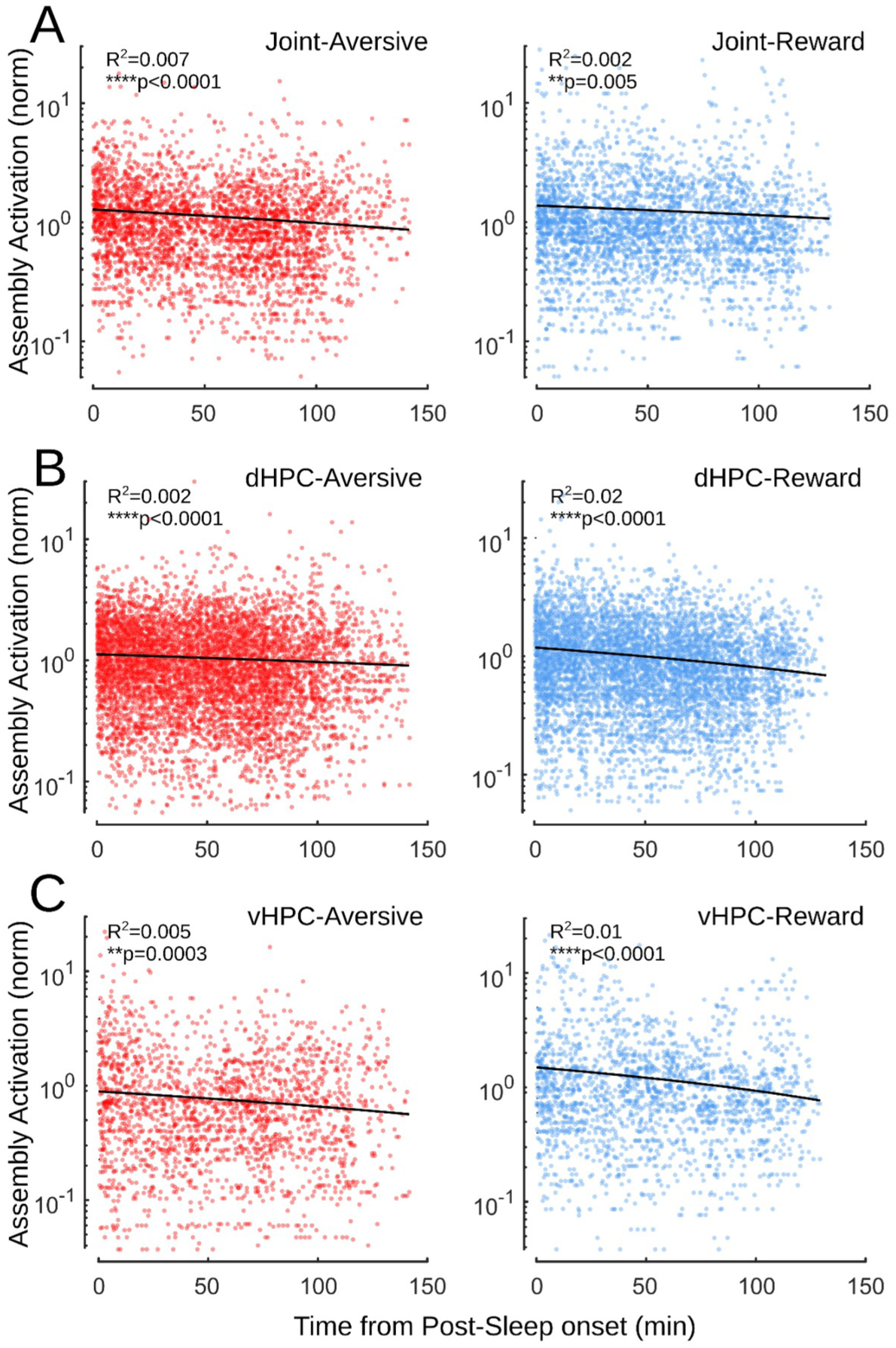
Dorsal-ventral hippocampal assembly activations decay over Post-NREM Sleep. **A**, Normalized Activation Rates of Joint aversive (red) and reward (blue) assemblies aligned to the beginning of the first post-NREM epoch (Linear regression, Aversive assemblies, R^2^=0.007, ****p_slope_<0.0001, N=3622; Reward assemblies, R^2^=0.002, **p_slope_=0.005, N=3346). **B**, Normalized Activation Rates of dorsal aversive and reward assemblies aligned to the beginning of the first post-NREM epoch (Linear regression, Aversive assemblies, R^2^=0.002, ****p_slope_<0.0001, N=8305; Reward assemblies, R^2^=0.02, ****p_slope_<0.0001, N=7389). **C**, Normalized Activation Rates of ventral aversive and reward assemblies aligned to the beginning of the first post-NREM epoch (Linear regression, Aversive assemblies, R^2^=0.005, **p_slope_=0.0003, N=2434; Reward assemblies, R^2^=0.01, ****p_slope_<0.0001, N=2364).

**Extended Data Figure 13:**
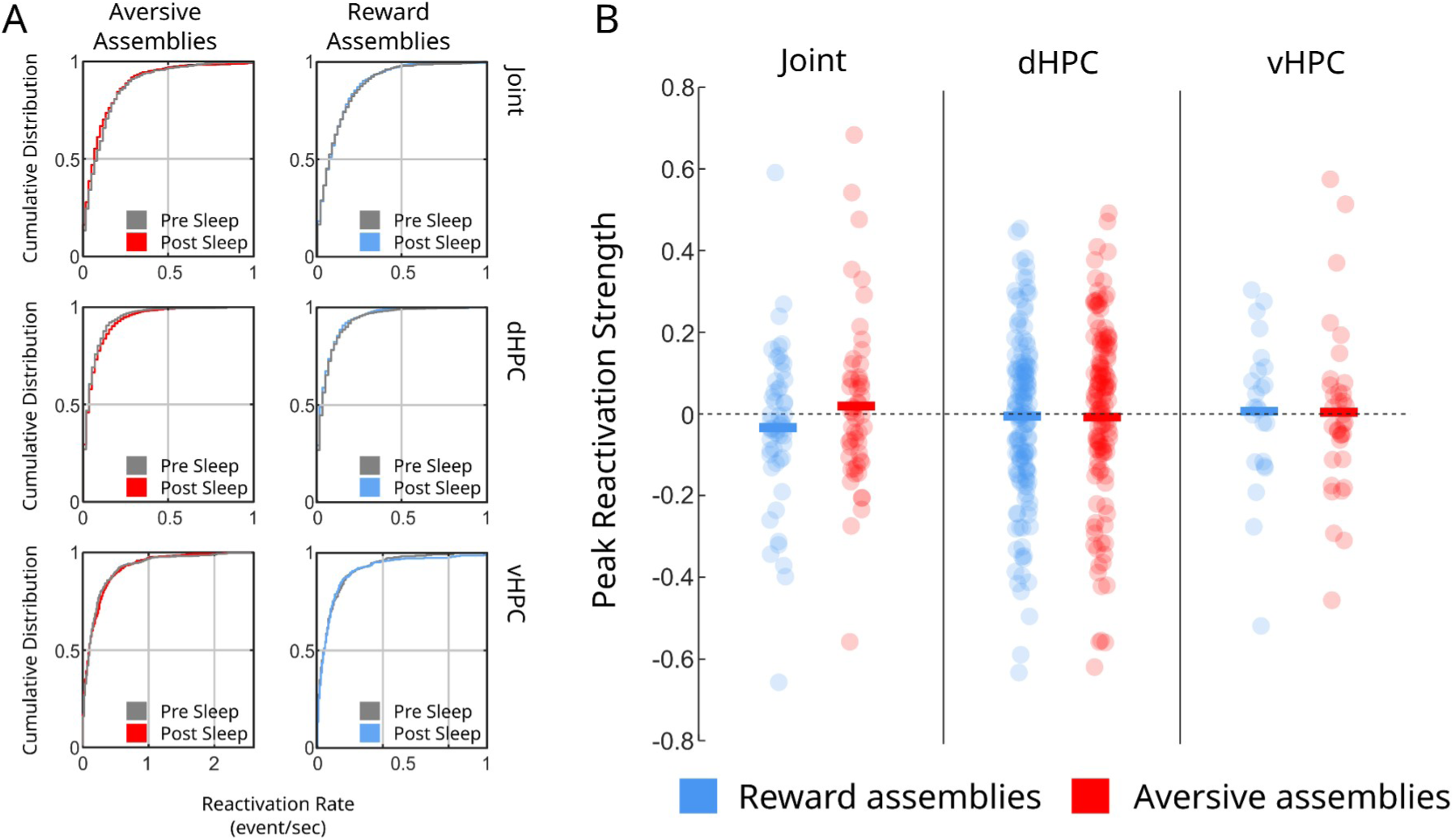
Assembly reactivation during REM sleep. **A**, Cumulative distribution of aversive and rewarded assembly reactivation rate in 60s bins during pre and post- NREM sleep for joint (top, dHPC (middle) and vHPC (bottom) assemblies. (Kolmogórov-Smirnov tests, p>0.05). **B**, Peak Reactivation Strength for all assembly types. (two-sided Mann-Whitney test. Joint: p=0.22, N_avesive_=54, N_reward_=53; dHPC: p=0.13, N_aversive_=131, N_reward_=146; vHPC: p=0.75, N_aversive_=37, N_reward_=34). We also compared the Peak Reactivation Strength against zero to infer increase of Reactivation Strength during Post-sleep compared Pre-sleep. (one-sided Mann-Whitney test against zero. Joint: p_reward_=0.86, p_aversive_=0.45; dHPC: p_reward_=0.47, p_aversive_=0.07; vHPC: p_reward_=0.21, p_aversive_=0.50)

**Extended Data Figure 14:**
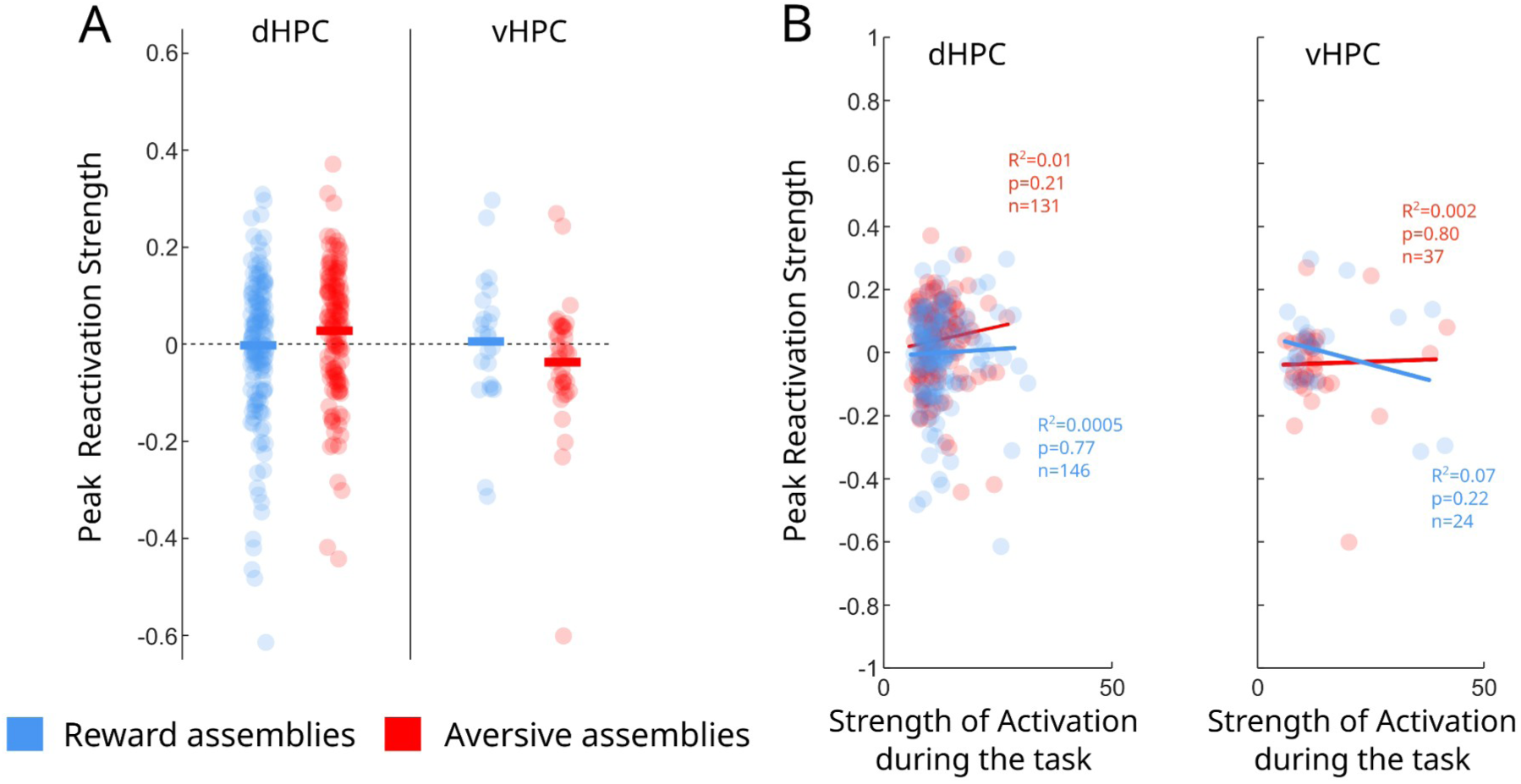
Peak Reactivation Strength for aversive and rewarded dorsal and ventral assemblies. **A**, We computed the Peak Reactivation Strength for reward (blue) and aversive (red) assemblies with members coming only from the dorsal (dHPC) or ventral hippocampus (vHPC; one-sided Mann-Whitney test against zero, p>0.05). **B**, We performed a correlation between the Peak reactivation Strength and the Activation Strength during the task for both type of assemblies showing members from the dHPC or vHPC. Linear correlation between the Peak Reactivation Strength and Activation Strength during the task. (Linear regression, Aversive assemblies, dHPC: R^2^=0.01, p_slope_=0.21, N_dHPC_=131, vHPC: R^2^=0.002, p_slope_=0.80, N_vHPC_=37; Reward assemblies, dHPC: R^2^=0.0005, p_slope_=0.77, N_dHPC_ =146, vHPC: R^2^=0.07, p_slope_=0.22, N_vHPC_=24).

**Extended Data Figure 15:**
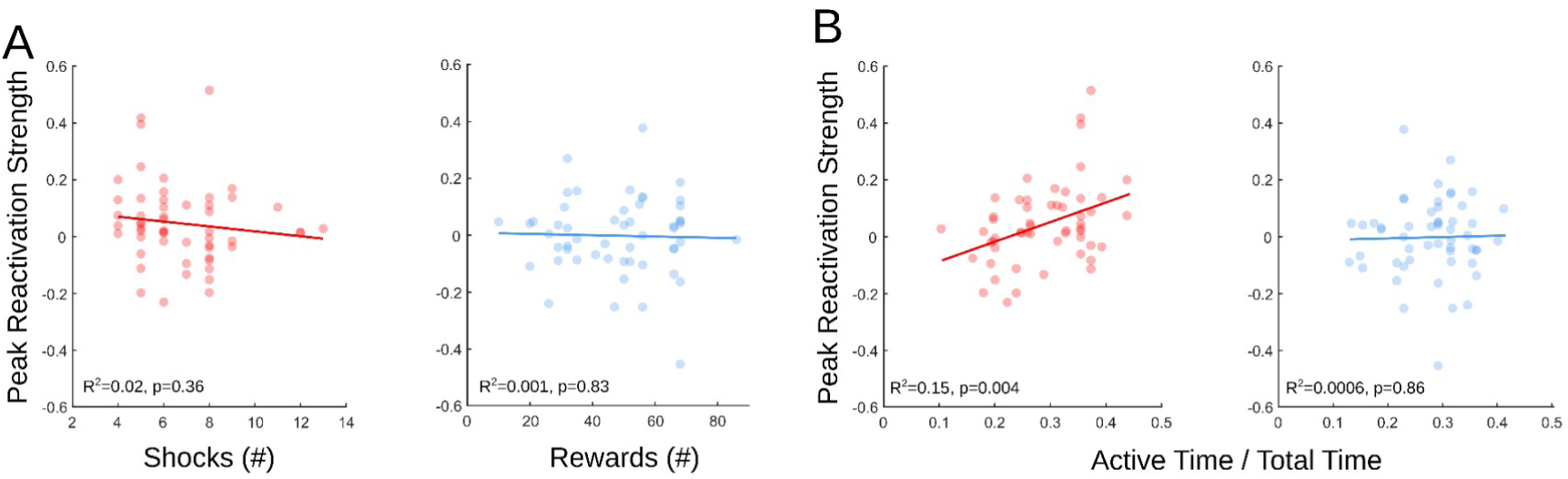
Peak Reactivation Strength correlates with Active time in Aversive run sessions. **A**, Correlation between Peak Reactivation Strength and Number of shocks (red) and number or rewards (blue) for aversive and reward joint-assemblies respectively (Linear regression, Aversive assemblies, R^2^=0.02, p_slope_=0.36, N=54; Reward assemblies, R^2^=0.001, p_slope_=0.83, N=53). **B**, Correlation between Peak Reactivation Strength Active Time during aversive (red) and reward (blue) run sessions for aversive and reward joint-assemblies respectively (Linear regression, Aversive assemblies, R^2^=0.15, **p_slope_=0.004; Reward assemblies, R^2^=0.0006, p_slope_=0.86).

**Extended Data Figure 16:**
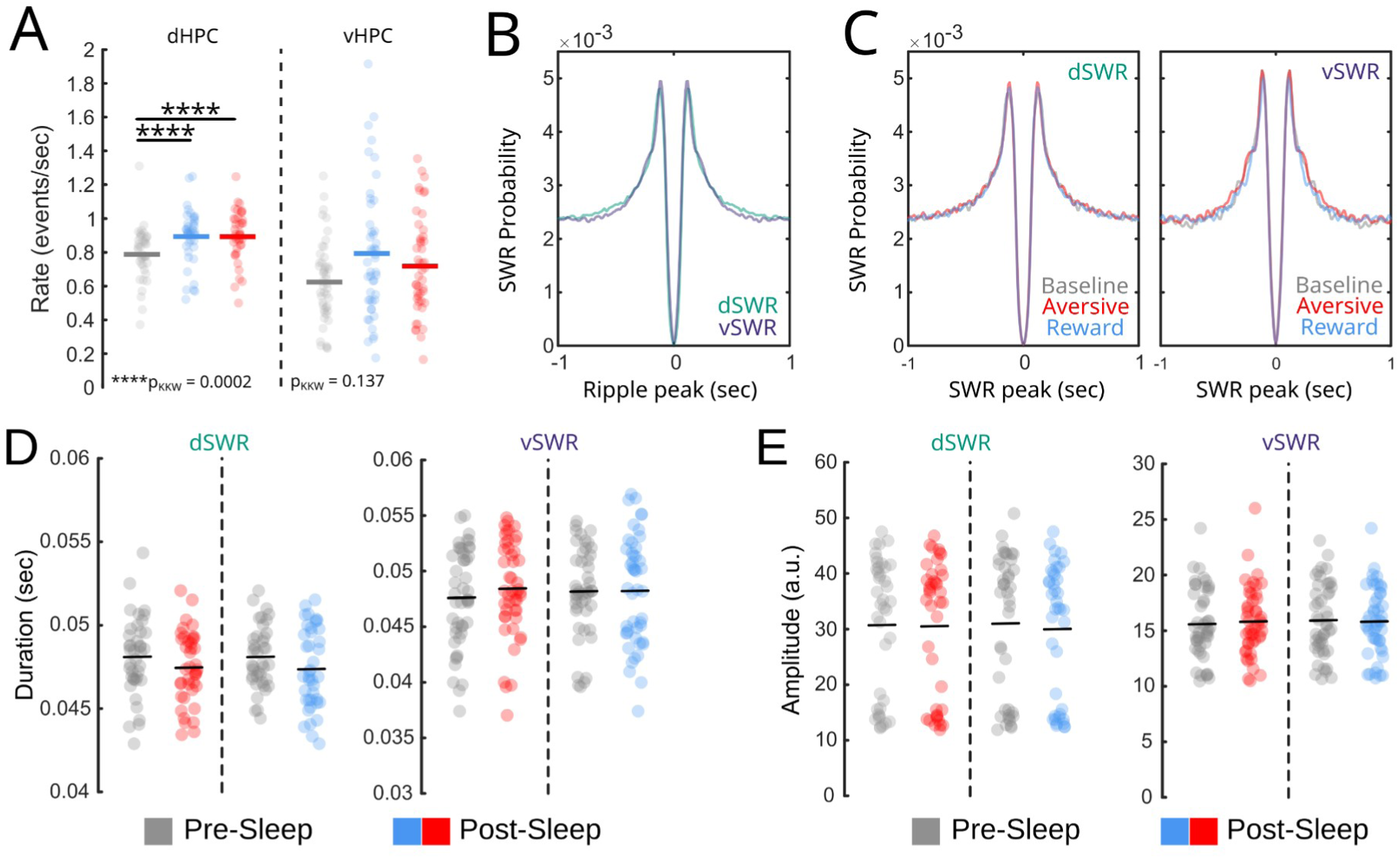
Characterization of dorsal and ventral ripples across conditions. **A**, Ripple rate in the dHPC and vHPC during NREM sleep in baseline (grey) and after the aversive (red) or rewarded (blue) condition. (Kruskal-Wallis test, ****p<0.0001, post-hoc test ****p<0.0001, N_dHPC_=40; N_vHPC_=47) detected in all sleep epochs. **B**, Autocorrelograms of dorsal and ventral ripple peaks during all NREM sleep, and **C**, separated in baseline, aversive, or reward NREM sleep. **D** _Mean duration of dorsal (dSWR) and ventral (vSWR) ripples detected in pre-sleep (gray) and post-sleep (red, aversive; blue, reward) sessions. (Paired Wilcoxon tests showed no significant differences. dSWR: paversive=0.21, preward=0.16; vSWR: paversive=0.35, preward=0.95). E, Mean Amplitude of dSWR and vSWR detected in pre-sleep and post-sleep sessions. (dSWR: paversive=0.84, preward=0.49; vSWR: paversive=0.79, preward=0.92. Ndorsal =40; Nventral =47)._

**Extended Data Figure 17:**
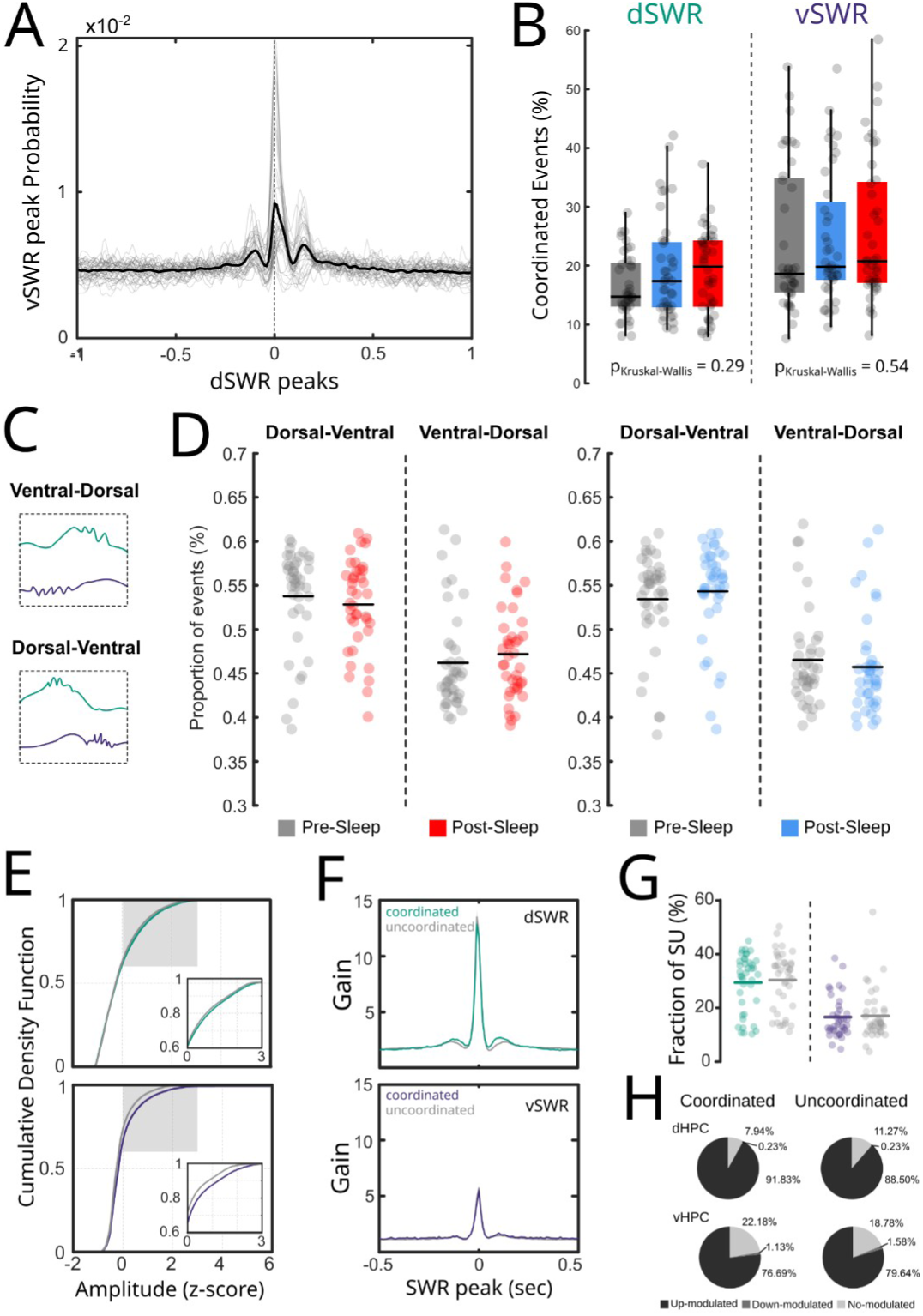
Characterization of dorsal-ventral ripple coordination across conditions. **A**, Peri-event density of ventral ripple peak density centered on dorsal ripple peaks for individual sessions (grey) and averaged across all sessions (black) showing the co-occurrence of dSWRs and vSWRs within a +/- 100ms window (dashed lines). **B**, Percentage of coordinated events across sleep sessions. (Kruskal-Wallis test, dSWRs: p=0.29; vSWRs: p=0.54, the line represents the median). **C-D** Percentage of ventral- and dorsal-leading events were segregated across pre- (in gray) and post-sleep (aversive in red, reward in light blue) sessions. (Paired Wilcoxon test; aversive, p = 0.19; reward, p = 0.27; N = 40). **E**, Cumulative density function of ripple peak amplitude for coordinated (green/purple) and uncoordinated ripples (grey). (Kolmogórov-Smirnov test, dSWRs, ****p<0.0001; vSWRs, ****p<0.0001). **F**, Peri-event activity gain for pyr single units for coordinated and uncoordinated dorsal (left) and ventral (right) ripples. **G**, Average fraction of single-units recruited during coordinated and uncoordinated ripples. (paired Wilcoxon test, p_dHPC_=0.18, , p_dHPC_=0.95, N=40). **H**, Percentages of modulated vs. non-modulated neurons for each ripple type. (Chi^2^ across structures: X^2^=6.35, p=0.1).

**Extended Data Figure 18:**
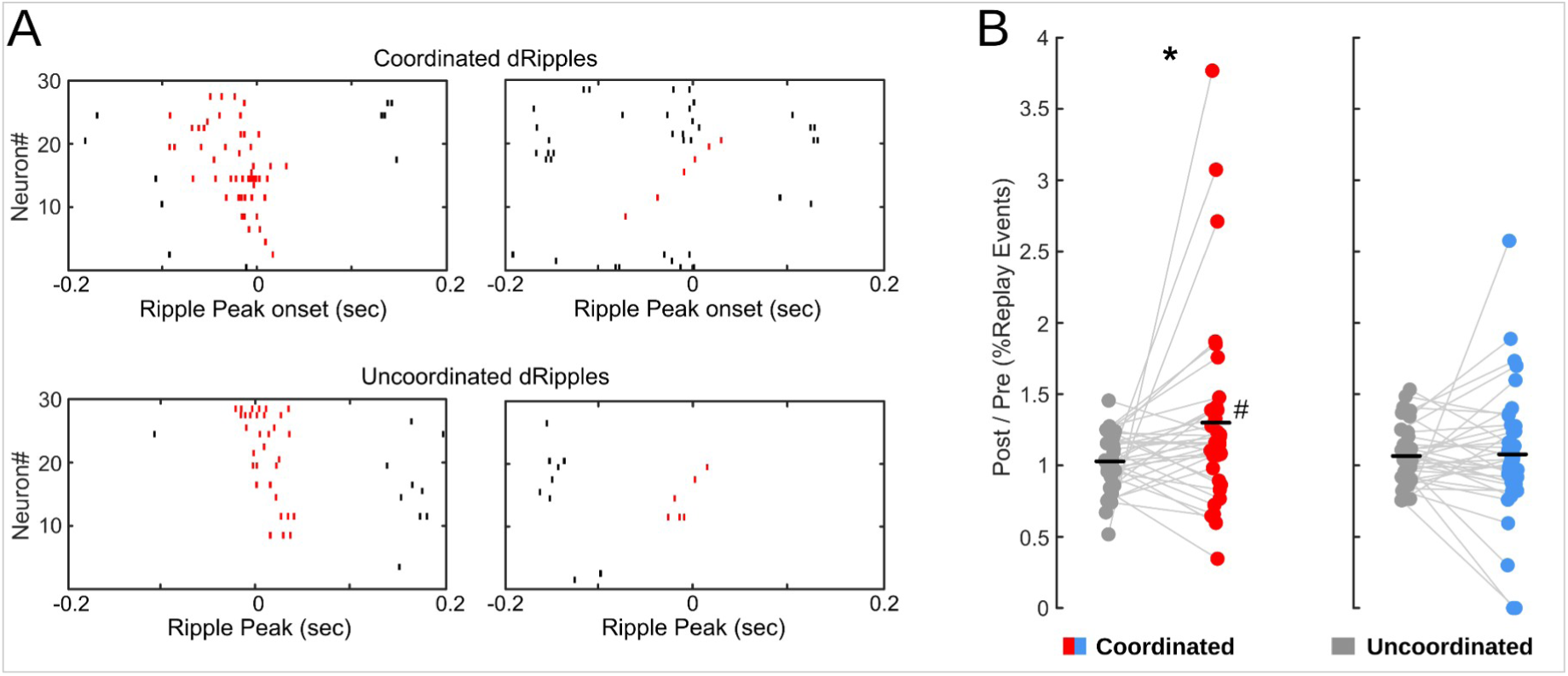
Dorsal replays detection using Rank-order method. **A**, Examples of significant dorsal replay events during dorsal ripples occurring within coordinated (top) and uncoordinated (bottom) ripple events. **B**, Replay ratio for coordinated (red or blue) and uncoordinated (gray) dorsal ripples. The percentage of significant replays detected during post-sleep was normalized to pre-sleep levels. Paired Wilcoxon test, (*p_aversive_=0.04, p_reward_=0.88, N=40. One-sample Wilcoxon test against 1, ^#^p=0.014).

**Extended Data Figure 19:**
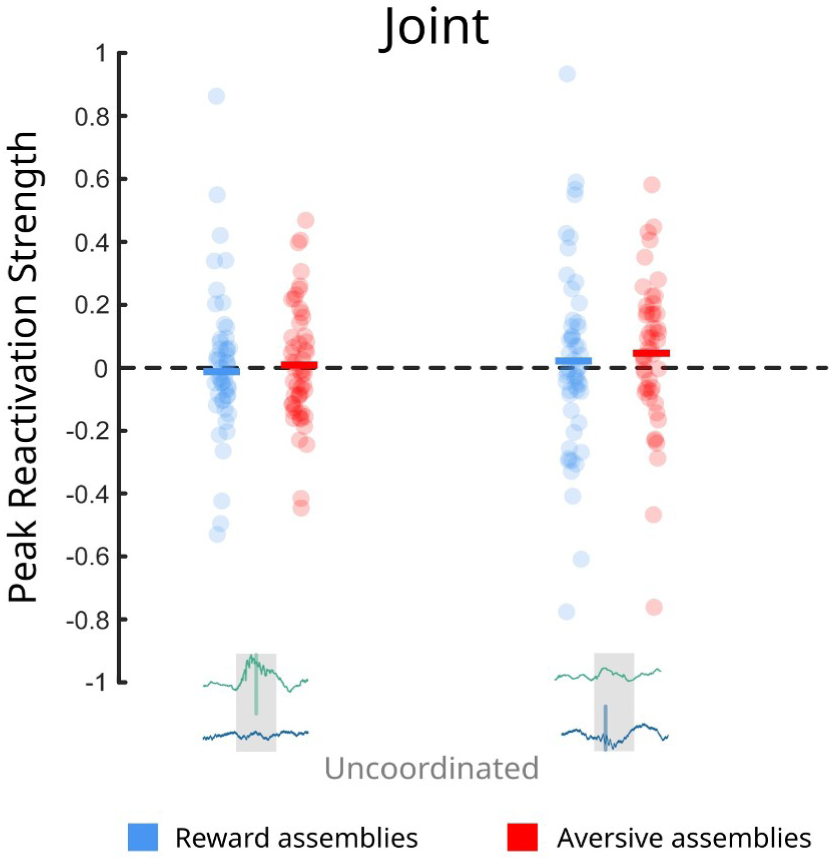
Peak Reactivation Strength of joint assemblies during dorsal and ventral uncoordinated events. We separated the Peak Reactivation Strength values coming from uncoordinated dorsal (left) and uncoordinated ventral ripples (right). (non-paired Two-way ANOVA, p_interaction_=0.91, F=0.01,p_assembly_type_=0.49, F=0.46, p_ripple_type_=0.46, F=0.54; one-sided t test test against zero, p>0.05).

**Extended Data Figure 20:**
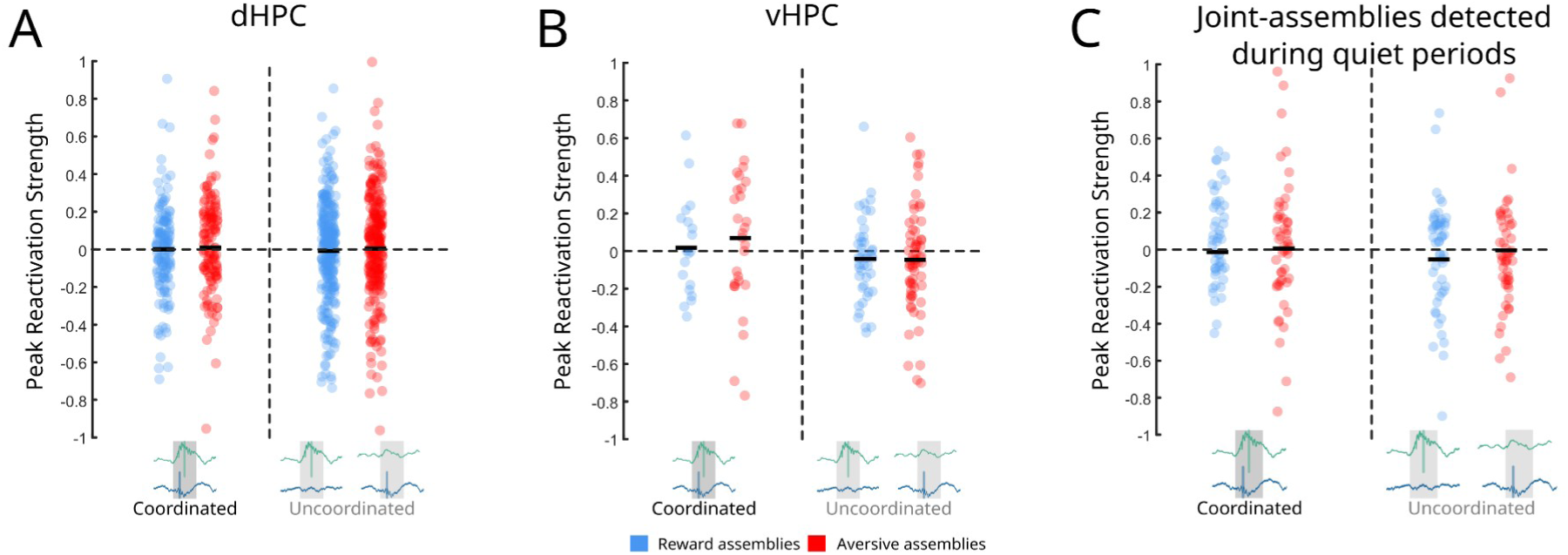
Peak Reactivation Strength of dorsal and ventral assemblies during coordinated and uncoordinated ripples. We compared Peak Reactivation Strength during coordinated and uncoordinated ripples, pooling dorsal and ventral uncoordinated events. **A**, Peak Reactivation Strength of dorsal aversive (red) and reward (blue) assemblies. (non-paired Two-way ANOVA, p_interaction_=0.95, F=0.004, p_assembly_type_=0.62, F=0.24, p_ripple_type_=0.70, F=0.15, N_aversive_=126, N_reward_=137). **B**, Peak Reactivation Strength of ventral aversive and reward assemblies. (non-paired Two-way ANOVA, p_interaction_=0.58, F=0.30, p_assembly_type_=0.66, F=0.19, p_ripple_type_=0.08, F=3.08, N_aversive_=29, N_reward_=19). **C**, Peak Reactivation Strength of joint reward- and aversive-assemblies detected during quiet periods of time. (Two-way ANOVA: p_interaction_=0.84, p_assembly_type_=0.25, p_ripple_type_=0.18, N_aversive_=57, N_reward_=57)

**Extended Data Figure 21:**
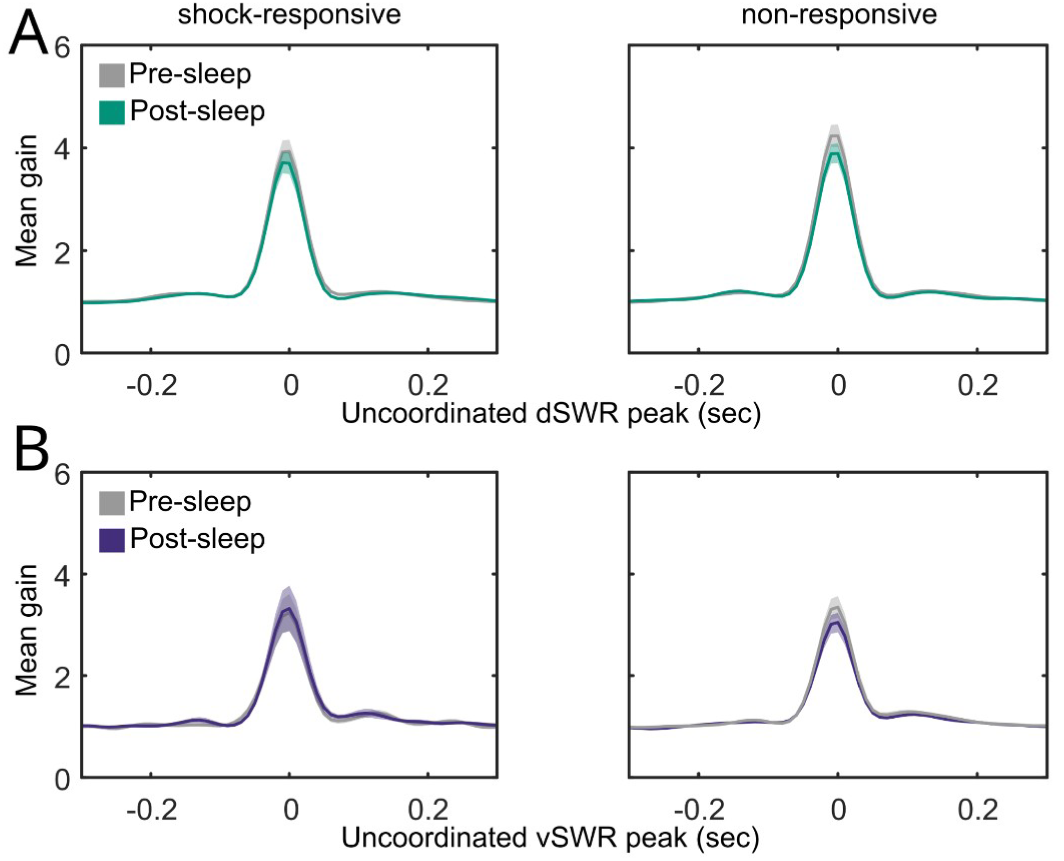
Response of shock- and non-responsive neurons to uncoordinated ripples. Mean peri- ripple gain for uncoordinated ripple onset in pre-and post-sleep of an aversive run in dHPC (**A**) and in vHPC (**B**). (one-sided Mann-Whitney test. dHPC, p_responsive_=0.99, n=84, p_non-responsive_=0.99, n=95. vHPC, p_responsive_= 0.22, n=53, p_non-responsive_=0.99 , n=120).

**Extended Data Figure 22:**
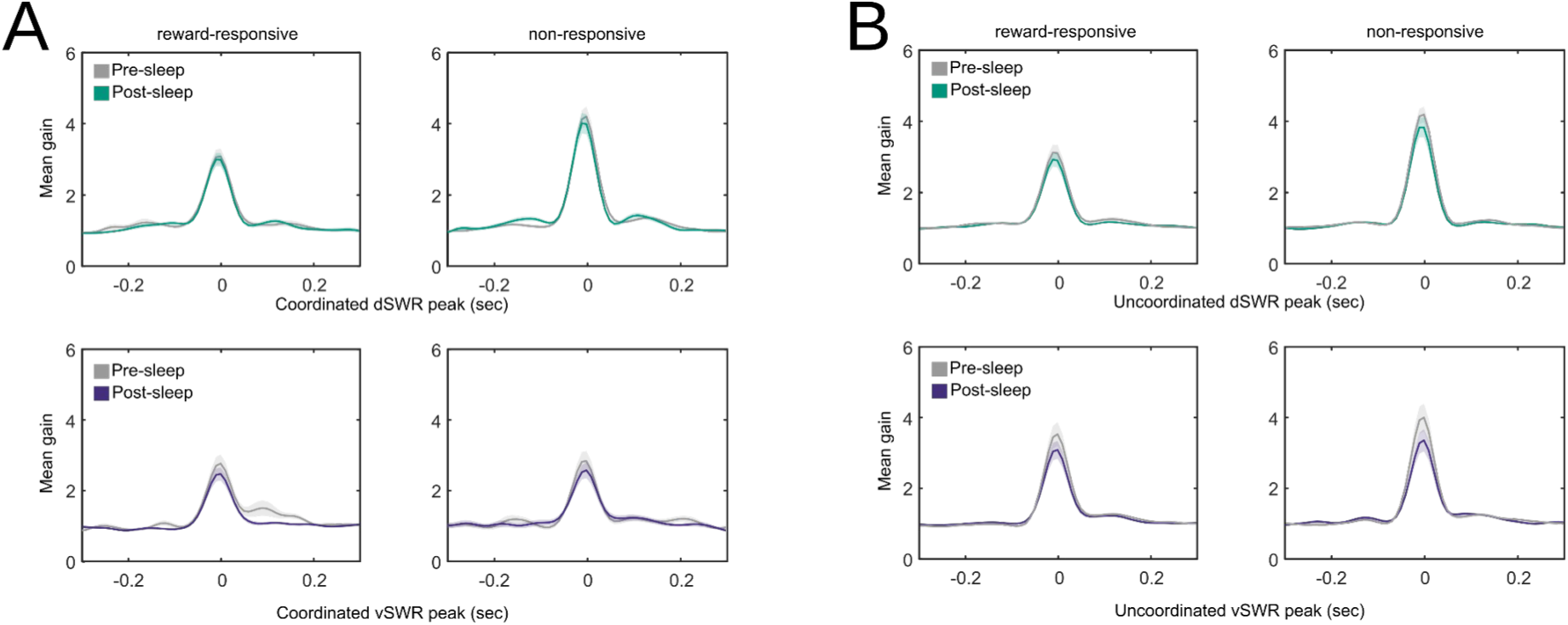
Reward-responsive Members from joint assemblies do not change their response during ripples. Reward-responsive cells response locked to coordinated and uncoordinated ripples. **A**, Average gain in firing rate around coordinated ripple onset during NREM sleep preceding (grey) and following (green/purple) reward runs. (one-sided Mann-Whitney test. dHPC, p_responsive_ =0.53, p_non-responsive_=0.80). (vHPC, p_responsive_=0.44 , p_non-responsive_=0.92). **B**, Average gain in firing rate around uncoordinated ripple onset during NREM sleep preceding (grey) and following (green/purple) reward runs. one-sided Mann-Whitney test. dHPC, p_responsive_=0.72, p_non-responsive_=0.87, N=62. vHPC, p_responsive_=0.43, p_non-responsive_=0.14, N=54.

**Extended Data Figure 23:**
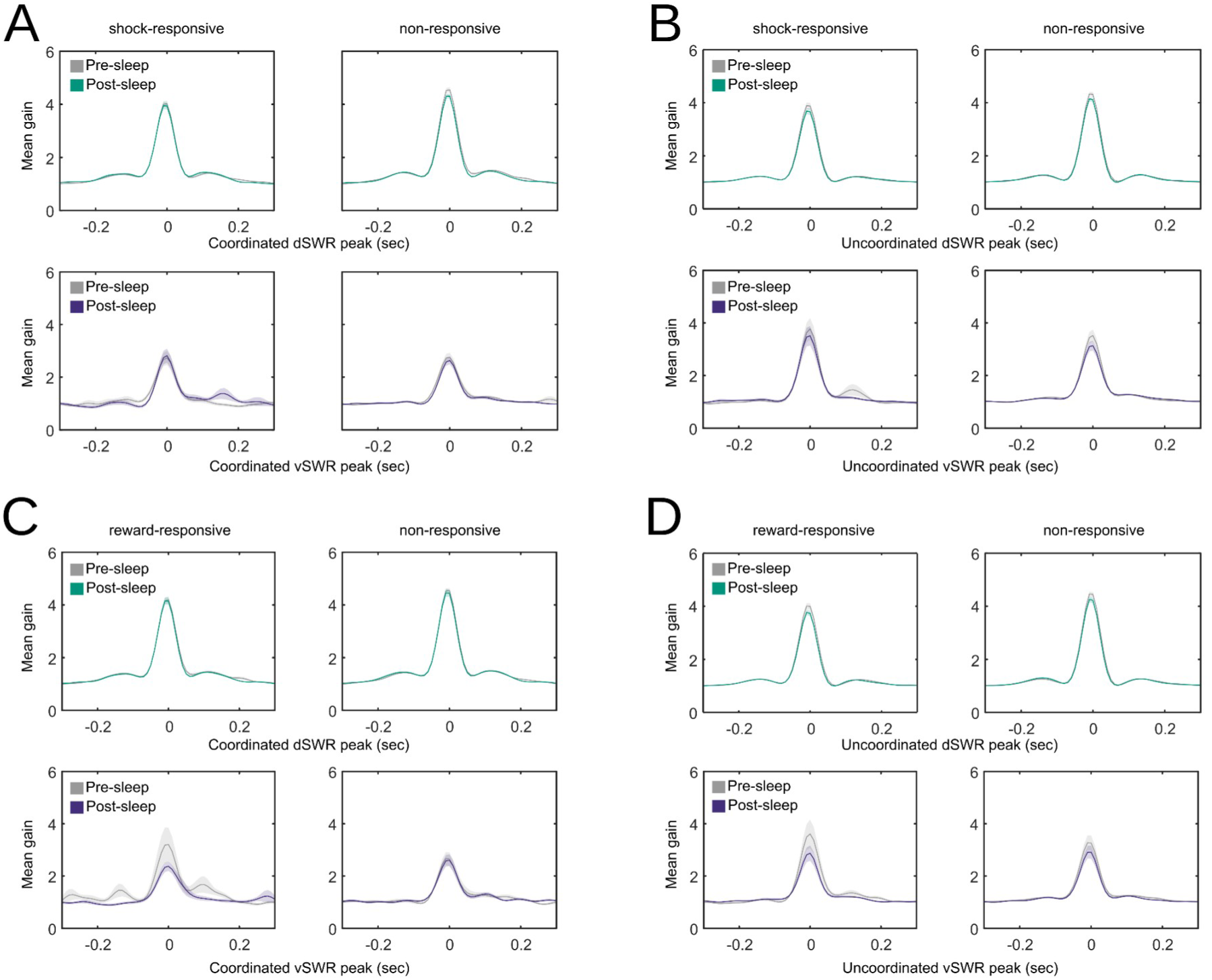
Members from purely dorsal or ventral assemblies do not change their response during ripples. Responses of shock- and reward-responsive versus non-responsive members from purely dorsal (upper panels) and purely ventral (lower panels) assemblies were tested with a one-sided Wilcoxon test. **A**, Shock-responsive cells during coordinated events. (dHPC responsive, p=0.91, N=200; vHPC responsive, p=0.88, N=53; dHPC non-responsive, p=0.99, N=200; vHPC non-responsive, p=98, N=53). **B**, Shock-responsive cells during uncoordinated events. (dHPC responsive, p=0.99, N=200; vHPC responsive, p=0.70, N=53; dHPC non-responsive, p=0.99, N=200; vHPC non-responsive, p=0.99, N=53). **C**, Reward-responsive cells during coordinated events. (dHPC responsive, p=0.98, N=227; vHPC responsive, p=0.13, N=49; dHPC non-responsive, p=0.99, N=227; vHPC non-responsive, p=0.98, N=49). **D**, Reward-responsive cells during uncoordinated events. (dHPC responsive, p=0.99, N=227; vHPC responsive, p=0.33, N=227; dHPC non-responsive, p=0.95, N=357; vHPC non-responsive, p=0.34, N=49).

